# Functional connectivity profile of the amygdala subfields associates with emotional well-being in aging

**DOI:** 10.64898/2026.02.04.703682

**Authors:** Shuer Ye, Arjun Dave, Xiaqing Lan, Menno P. Witter, Alireza Salami, Maryam Ziaei

**Author notes:** Correspondence should be addressed to Maryam Ziaei, or Shuer Ye.

## Abstract

Amygdala-related functional connectivity plays a crucial role in human emotion, cognition, and mental well-being. The amygdala is a highly heterogeneous structure, with subregions that have both distinct and overlapping functions. However, the connectivity patterns of different amygdala subregions—and how these associations vary with age—remain poorly understood.

Functional MRI data were analyzed from 68 younger adults and 66 older adults during movie watching in a 7T MRI scanner. Partial least squares (PLS) analysis was used to identify latent variables capturing variance associated with age-related differences in the functional connectivity patterns of three amygdala subregions: the basolateral (BLA), centromedial (CMA), and superficial (SFA) nuclei. In addition, covariance between behavioral measures, such as emotional resilience and cognitive function, and functional connectivity of these subregions was examined. We further explored the associated cognitive processes, correspondence with large-scale brain networks, and the underlying chemoarchitecture of the identified connectivity patterns of the BLA, CMA, and SFA.

Multivariate analyses revealed age-related and subregion-specific functional connectivity patterns. Functional connectivity patterns of amygdala subregions were further associated with emotional resilience, which largely overlapped across widespread brain regions; however, their associations differed by age. Stronger coupling of amygdala subregions predicted higher resilience in older adults but lower resilience in younger adults. The identified connectivity patterns were linked to the salience, control, and default mode networks and showed spatial correspondence with mGluR5 and 5-HT6 receptor distributions.

These findings highlight age-dependent reorganization of amygdala-related networks that support emotional resilience and provide novel insights into how functional and neurochemical changes of the amygdala subregions contribute to adaptive emotional and cognitive functions in aging.

## Introduction

The amygdala is a central hub of emotional processing and a key node of the salience network [SN], maintaining widespread connections with other brain regions to support emotional and cognitive functions (Pessoa, 2010; Phelps & LeDoux, 2005; Seeley, 2019a). Structurally, the amygdala is a heterogeneous structure comprising the basolateral (BLA), centromedial (CMA), and superficial (SFA) subregions, each exhibiting distinct yet partially overlapping connectivity profiles. The BLA acts as an “input hub,” integrating cortical, thalamic, and hippocampal information to evaluate the emotional significance of stimuli (Janak & Tye, 2015; Sladky et al., 2013), whereas the CMA functions as the principal “output hub,” transmitting affective evaluations to hypothalamic, brainstem, and striatal regions to orchestrate autonomic and behavioral responses (Allen et al., 2021; Dall’Oglio et al., 2015). The SFA is primarily involved in olfactory and social-affective processing through its connections with the insula, ventral striatum, and other limbic areas (Heimer & Van Hoesen, 2006; Price, 2003). Functional neuroimaging and connectivity studies consistently demonstrate subregional specialization: the BLA connects with prefrontal–parietal control network (CN), the CMA with motor and striatal systems, and the SFA with limbic circuits (Alarcón et al., 2015; Bzdok et al., 2013; Kirstein et al., 2023). These connectional distinctions likely associate with behavioral and clinical functionality, as aberrant subregional connectivity has been implicated in depression, obsessive–compulsive disorder, and Parkinson’s disease (Jacob et al., 2022; Kwon et al., 2024; Wang et al., 2023).

As the brain ages, the amygdala and its subregions undergo morphological changes. Most volumetric MRI studies have treated the amygdala as a single structure, reporting an overall age-related volume decline (Fjell et al., 2013; Raz et al., 2003; Walhovd et al., 2011). However, emerging evidence points to heterogeneous, nonlinear trajectories across amygdala nuclei, with the BLA showing negative associations with age whereas the CMA exhibits relative stability, potentially reflecting differences in function, development, or genophylogenetic origin (Aghamohammadi-Sereshki et al., 2019; Kurth et al., 2019). Age-related cellular and molecular alterations, including centromedial neuronal loss and amyloid accumulation in cortical transition zones, have been linked to Alzheimer’s pathology (Nikolenko et al., 2020). Notably, volumetric reductions across multiple amygdala subfields have been observed in individuals with mild cognitive impairment despite preserved global gray matter (Singh et al., 2024), highlighting the amygdala’s vulnerability during aging and neurodegeneration. Although structural changes in the amygdala and its subregions have been widely linked to aging, much less is known about how aging affects its functional coupling with the rest of the brain.

Previous research on large-scale brain networks has revealed age-related functional dedifferentiation characterized by reduced network segregation and increased integration (Chan et al., 2014; Damoiseaux, 2017). Such large-scale reorganization has been linked to changes in cognitive performance and emotional regulation efficiency in late adulthood (Deery et al., 2023; Spreng & Turner, 2019; Ye et al., 2025). Given the amygdala’s central role in integrating affective and cognitive processes, functional alterations within its subregions may help explain the emotional changes observed in older adults, such as enhanced emotional stability and a selective focus on positive information (Carstensen et al., 2011; Isaacowitz et al., 2006; Ziaei et al., 2017). Empirical evidence suggested that older adults show stronger functional connectivity between the amygdala and the default mode network (DMN) following negative emotional experiences (Baez-Lugo et al., 2023). Additionally, stronger functional coupling between the prefrontal cortex and the amygdala has been linked to reduced negative affective ratings in older adults, providing a neural account for the positivity bias commonly observed in aging (Sakaki et al., 2013; St. Jacques et al., 2010). Despite these insights, research investigating how the functional coupling of amygdala subregions relates to emotional well-being and cognitive function, and how these relationships are modulated by aging, remains scarce. Understanding how amygdala-related networks support emotional and cognitive changes in later life may guide strategies to enhance psychological well-being and cognitive health in aging.

To address this gap, the present study used ultra–high-field 7T MRI, which offers superior spatial resolution for gaining the precise functional response of small nuclei within the amygdala (Qin & Gao, 2021; Sladky et al., 2013), combined with a naturalistic movie-watching paradigm that evokes dynamic, ecologically valid emotional responses (Eickhoff et al., 2020; Finn, 2021). Younger and older adults underwent a movie-watching fMRI session in the 7T MRI scanner, during which they were exposed to rich and dynamic stimuli featuring both negative and neutral content. Using multivariate partial least squares (PLS) analysis, we investigated how movie-watching functional connectivity profiles of individual amygdala subregions differ between two age groups, how the connectivity profile is related to emotional resilience and cognitive function based on age groups. Furthermore, to gain a more comprehensive understanding of these functional connectivity patterns and to assess distinctions among amygdala subregions, we examined their associations with cognitive processes, large-scale network correspondence, and neurochemical correlates.

Building on prior evidence demonstrating the amygdala’s role as a central hub for emotional processing and its associated functional connectivity, we expected that a multivariate method would identify unique and shared networks related to the three amygdala subregions, as well as age-related differences. We also anticipated that the connectivity patterns of different subregions would be related to behavioral measures, particularly emotional resilience. Finally, we anticipated that the identified functional patterns would involve large-scale systems known to support affective and regulatory processes (e.g., DMN, CN, and SN), although they may differ in their specific engagement patterns (Bzdok et al., 2013; Spreng & Turner, 2019; Ye et al., 2025). Moreover, given the evidence for age-related alterations in amygdala connectivity (Dolcos et al., 2020; Ziaei et al., 2017) and broader functional dedifferentiation (Baez-Lugo et al., 2023; Chan et al., 2014), we hypothesized that older adults would exhibit stronger functional coupling between the identified networks and amygdala subregions, whereas younger adults would show weaker coupling patterns compared to older counterparts for the amygdala networks.

## Method

### Participants

The current study analyzed data collected within the Trondheim Aging Brain Study (TABS), which investigates emotional processing during aging (Dave et al., 2025). Participants comprised 80 younger adults and 80 older adults, recruited from local communities through advertisements. All were healthy, had normal hearing, and either normal or corrected-to-normal vision, with no MRI contraindications or history of psychiatric or neurological conditions. After exclusions due to technical problems, incomplete datasets, or excessive movement (mean frame displacement [FD] > 0.3 mm), the final analysis included 68 younger adults (32 females, mean age 25.91 ± 4.31 years, range 19–36 years) and 66 older adults (34 females, mean age 71.02± 3.81 years, range 65–82 years). Participants attended two sessions—a behavioral and an imaging session—provided written informed consent and received a 500 NOK gift card as compensation. The research complied with the Declaration of Helsinki and had ethical approval from the regional committees for medical and health research ethics (REK-Midt, approval #390390).

### Task design

This study employed two distinct movie clips during functional MRI scanning: one neutral and one negative. The neutral clip, titled “Pottery,” shows two women engaged in making pottery, lasting for 480 seconds. In contrast, the negative clip, “Curve,” features a woman desperately trying to prevent herself from slipping off a cliff into an abyss, with a duration of 494 seconds. These clips were chosen based on results from a pilot study that evaluated emotional assessment, arousal, and valence ratings from four potential clips. The selected clips demonstrated strong validity in eliciting neutral and negative emotional states, thus being integrated into the imaging experiment. Additional details about these clips are available in our prior publication (Ye et al., 2024), where we confirmed that “Curve” effectively induces stress and emotional arousal, whereas “Pottery” functions as an appropriate neutral control, evoking minimal emotional reactions according to participants’ self-reports. Consequently, this study emphasizes the negative movie clip as the primary condition, using the neutral clip as a control.

### Neuropsychological assessments

Participants completed a comprehensive battery of neuropsychological and self-report measures administered outside the MRI environment to assess emotional well-being and cognitive performance. Emotional and psychological well-being were evaluated using validated questionnaires, including the Depression, Anxiety, and Stress Scale (Lovibond & Lovibond, 1995), Difficulties in Emotion Regulation Scale (Gratz & Roemer, 2004), General Health Questionnaire (Werneke et al., 2000), Hospital Anxiety and Depression Scale (Bjelland et al., 2002), Connor–Davidson Resilience Scale (Connor & Davidson, 2003), Intolerance of Uncertainty Scale (Buhr & Dugas, 2002), Perceived Stress Scale (Cohen et al., 1983), and the State–Trait Anxiety Inventory (Gunning et al., 2010). Cognitive function was assessed using standard tests, including the Stroop test (Jensen & Rohwer, 1966), forward and backward digit span tests (Schroeder et al., 2012), Trail Making Test Parts A and B (Reitan & Wolfson, 2004), and the Controlled Oral Word Association Test for phonemic and semantic verbal fluency (Ivnik et al., 1996). Empathy and theory of mind were evaluated using the Reading the Mind in the Eyes Test (Baron-Cohen et al., 2001) and the Interpersonal Reactivity Index (Davis, 1983), and general intelligence was assessed with the abbreviated Raven’s Progressive Matrices (Raven, 1941).

To reduce dimensionality and derive composite behavioral indices, Principal component analysis was performed separately on emotional and cognitive measures (Abdi & Williams, 2010). This yielded two primary components: the Emotional Resilience Index (ERI), representing emotional resilience and mental well-being, and the Cognitive Function Index (CFI), reflecting general cognitive performance. Sampling adequacy was high for the ERI (Kaiser–Meyer–Olkin [KMO] = 0.918; Bartlett’s test: χ²(142) = 105, p < 0.001) and moderate for the CFI (KMO = 0.67; Bartlett’s test: χ²(142) = 21, p < 0.001). Significant principal components had eigenvalues greater than 1. The ERI comprised two components explaining 63.94% of the total variance (57.05% and 6.89%), whereas the CFI included three components explaining 72.69% of the variance (37.26%, 20.06%, and 15.37%). Composite ERI and CFI scores were computed as weighted sums of factor scores based on the variance explained (Dave et al., 2025). Higher ERI scores indicate greater emotional resilience, whereas higher CFI scores reflect superior cognitive ability across verbal fluency, working memory, and inhibitory control domains.

### Image acquisition

Imaging acquisition was conducted using a Siemens MAGNETOM Terra 7T MRI scanner equipped with a 32-channel head coil at the Norwegian 7T MR Center, St. Olav Hospital. Functional images were acquired with a multi-band accelerated echo-planar imaging sequence (92 interleaved slices, multi-band acceleration factor = 2, voxel dimensions = 1.25 × 1.25 × 1.25 mm, TR = 2000 ms, TE = 19 ms, matrix size = 160 × 160 mm, field of view = 200 mm, slice thickness = 1.25 mm, flip angle = 80°), generating 243 volumes for the neutral movie and 250 volumes for the negative movie. Anatomical images were acquired using an MP2RAGE sequence with 224 sagittal slices (voxel dimensions = 0.8 × 0.8 × 0.8 mm, TE = 1.99 ms, TR = 4300 ms, inversion times = 840 ms and 2370 ms, flip angles = 5°/6°, slice thickness = 0.75 mm).

### Image preprocessing

The preprocessing were conducted using fMRIPrep version 22.0.2 (Esteban et al., 2019) and XCP-D version 0.3.053 (Mehta et al., 2024), respectively. Raw imaging data in DICOM format were converted to NIfTI format while maintaining original acquisition parameters. The first three volumes were discarded to eliminate initial transient MR signals. Functional data preprocessing followed the standard pipeline in fMRIPrep, which comprised slice timing correction, realignment, co-registration to the anatomical T1-weighted images using boundary-based registration (FreeSurfer; Greve and Fischl, 2009), and normalization to the MNI standard space. Pos-preprocessing, including nuisance regression, band-pass filtering, and spatial smoothing with 4mm kernel size, was then implemented with XCP-D pipeline. More details of preprocessing strategies can be found in **supplementary material**.

### Amygdala subregions

To accommodate the high-resolution 7T functional MRI images, we employed the Tian subcortical atlas (7T-SCALE3) (Tian et al., 2020). The atlas was generated by mapping resting-state functional gradients using Laplacian eigenmaps, with discrete subcortical boundaries delineated only when supported by model selection, while continuous gradient information was preserved elsewhere. Compared with traditional anatomy-based parcellations, the subcortical atlas derived from functional gradients captures continuous topographic organization while providing reproducible discrete subdivisions that align with cortical networks, thereby offering a more functionally informed and behaviorally relevant characterization of the human subcortex. In this atlas, the amygdala is parcellated into three subregions, corresponding to the bilateral BLA, CMA, and SFA (**Fig.S1**).

We performed tissue segmentation and volume estimation to volume of the amygdala and its subregions by using the CAT12 toolbox based on MATLAB2023b (Gaser et al., 2024). T1-weighted structural images were first preprocessed using affine regularization, correction for bias-field inhomogeneity, and spatial normalization to the Montreal Neurological Institute (MNI) template. The modulated gray matter maps were smoothed using an 8□mm full-width at half-maximum (FWHM) Gaussian kernel. Based on the amygdala parcellation from Tian’s atlas (Tian et al., 2020), the gray matter (GM) maps derived from tissue segmentation were masked by these subregion templates to extract the corresponding GM volumes. The mean modulated GM value within each subregion was then summed to obtain the total volume of that subregion for each participant. During preprocessing, total intracranial volume (TIV), that is the overall volume, in cubic millimeters, of the space within the skull, was calculated for each subject. The subregional volumes and TIV were included as covariates in subsequent analyses. Notably, TIV did not differ significantly between groups (*t* = −1.64, *p* = 0.102, *d* = −0.28). However, older adults had smaller whole amygdala volume (*t* = 5.81, *p* < 0.001, *d* = 1.00), as well as reduced SFA (*t* = 5.87, *p* < 0.001, *d* = 1.12), BLA (*t* = 5.06, *p* < 0.001, *d* = 0.87), and CMA (*t* = 6.71, *p* < 0.001, *d* = 1.16) subregion volumes compared with younger adults after correction for the TIV.

### Seed-based amygdala FC

We estimated amygdala FC using three seed regions. Because the current study was not focused on lateralized effects, the left and right amygdala subregions were combined into bilateral masks, resulting in three ROIs: the SFA, CMA, and BLA. For each ROI, the mean BOLD signal was extracted by averaging across all voxels within the ROI, for both left and right side, and whole-brain FC was then computed by correlating this time series with the time series of every voxel in the brain using Pearson correlation.

### Partial least squared analyses

Following our previous work (Ziaei et al., 2016, 2021, 2022), we applied Partial Least Squares (PLS) analysis (Krishnan et al., 2011), a data-driven multivariate technique, to investigate (a) amygdala connectivity patterns by examining unique and shared coupling across age groups and subregions, and (b) age-related functional connectivity patterns in relation to emotional resilience and cognitive reserve. In contrast to traditional univariate approaches, this multivariate framework evaluates all behavioral measures and groups simultaneously, allowing for the identification of data-driven patterns of shared variance without the need for multiple-comparison correction (Zöller et al., 2017).

PLS uses singular value decomposition (SVD) on a cross-correlation matrix constructed by relating amygdala subregion FC with behavioral measures across participants, yielding a set of orthogonal latent variables (LVs). The first LV typically accounts for the largest proportion of shared covariance, with subsequent LVs explaining progressively smaller proportions. Each LV represents a whole-brain FC pattern (voxel saliences) that is optimally associated with behavioral measures, together with a singular value indicating the proportion of covariance explained. The statistical significance of each LV was evaluated via permutation testing with 5000 random permutations (McIntosh et al., 1996). For significant LVs, Pearson correlations between brain scores from significant LVs and behavioral measures were further examined to characterize the interplay between brain function, age groups, and behavior. Participants’ individual contributions to each LV were expressed as brain scores, obtained by projecting each participant’s imaging data onto the voxel saliences of a given LV (i.e., dot product). These scores correspond to the left singular vectors from the SVD and indicate how strongly an individual expresses the identified brain pattern and brain–behavior pattern.

The reliability of voxel saliences was assessed using bootstrap estimation, with confidence intervals derived from 500 bootstrap resamples (Efron & Tibshirani, 1985); voxel saliences with bootstrap ratios (BSR = loading/SE) exceeding |3.3| were considered reliable (approximately *p* < 0.001; Sampson et al., 1989; Wiebels et al., 2016), and a minimum cluster extent of 400 mm³ (equivalent to ≈205 voxels at 1.25-mm isotropic resolution) was further applied to reduce spurious small clusters. The reliability of behavioral loadings was assessed using the same bootstrap procedure; 95% confidence intervals (CIs) were computed, and loadings whose CIs did not include zero were deemed significant, indicating a reliable contribution of the variable to the corresponding LV.

Before conducting the PLS analyses, the effects of sex, head motion, TIV, and the volume of corresponding amygdala subregions were regressed out from the FC maps at the voxel-wise level. We then performed PLS analyses on the whole-brain functional connectivity patterns of the three amygdala subregions (SFA, CMA, and BLA) to further characterize their similarities and differences in age-related effects and further associations with behavioral measures.

### Network correspondence analysis

To investigate the associations between the salience maps and canonical functional networks (e.g., the salience and default mode networks), network correspondence analyses were performed using the Network Correspondence Toolbox (R. Kong et al., 2025) implemented in Python 3.7. This analysis provided a quantitative evaluation of the spatial correspondence between the salience maps and canonical network templates. The overlap with established functional network atlases was quantified using Dice coefficients, with higher values indicating greater spatial overlap between the two maps. The statistical significance of the Dice coefficients was assessed using a spin test with 1,000 random rotations of the input salience maps. Two canonical brain atlases—the Gordon2017-17 and Yeo2011-17 parcellations, each comprising 17 spatially distinct functional networks—were used as reference templates (Gordon et al., 2017; Thomas Yeo et al., 2011). Details of two atlas can be found in **Fig.S2**

### Cognitive processes decoding

To better interpret the related function of salience maps, we assessed their correspondence with canonical cognitive maps to infer the potentially related cognitive processes (Hansen et al., 2024; Luppi et al., 2025). Probabilistic associations between brain voxels and cognitive processes were extracted using Neurosynth, a meta-analysis platform aggregating findings from over 14,000 published fMRI studies. Neurosynth identifies these associations by analyzing commonly occurring keywords (e.g., ‘pain’, ‘attention’) reported alongside fMRI activation coordinates, specifically utilizing volumetric association test maps. The resulting measure reflects the likelihood that a particular cognitive process is reported when activation is observed at a specific voxel. Among the more than 1,000 cognitive processes available in Neurosynth, our analysis focused on 123 cognitive and behavioral terms derived from the Cognitive Atlas (Poldrack et al., 2011)—an open, comprehensive ontology for cognitive science—covering broad domains (e.g., attention, emotion), specific cognitive functions (e.g., visual attention, episodic memory), behavioral activities (e.g., eating, sleep), and emotional states (e.g., fear, anxiety). (Schaefer et al., 2018). Neurosynth-derived voxel maps were parcellated into 200 regions using the Schaefer atlas to construct a 200 × 123 (regions × terms) matrix. The salience maps were then parcellated using the same template, and correlations were computed between the canonical cognitive maps and the salience maps to identify associated terms with their corresponding *r*-values. These *r*-values reflect the degree to which each cognitive term’s meta-analytic activation pattern (from Neurosynth; Yarkoni et al., 2011) aligns with the spatial distribution of the salience maps.

### Association with neurotransmitters density

To investigate the underlying chemoarchitectural basis of the salience maps, we examined their associations with neurotransmitter density maps derived from previous PET studies by using JuSpace v2.0 toolbox (Dukart et al., 2021). Nineteen neurotransmitter maps included in this study represented the following receptor and transporter systems: 5HT1A (serotonin receptor subtype 1A), 5HT1B (serotonin receptor subtype 1B), 5HT2A (serotonin receptor subtype 2A), 5HT4 (serotonin receptor subtype 4), 5HT6 (serotonin receptor subtype 6), D1 (dopamine D1 receptor), D2 (dopamine D2 receptor), DAT (dopamine transporter), NMDA (N-methyl-D-aspartate receptor), GABAa (gamma-aminobutyric acid receptor A), mGluR5 (metabotropic glutamate receptor type 5), CB1 (cannabinoid receptor 1), M1 (muscarinic acetylcholine receptor), VAChT (vesicular acetylcholine transporter), NAT (noradrenaline transporter), MOR (μ-opioid receptor), H3 (histamine H3 receptor), and 5HTT (serotonin transporter). The neurotransmitter density maps selected were the most used in literature and were derived from the largest number of healthy individuals (Hansen et al., 2022). Salience maps were used as inputs for spatial correlation analyses with neurotransmitter maps. The default Neuromorphometrics atlas, provided in Juspace, which excludes all white matter and cerebrospinal fluid regions, was used to extract mean values for each region from both the input salience maps and the selected neurotransmitter/metabolic maps (Friston et al., 1994). Spearman correlation coefficients were then computed to quantify the spatial correspondence between the salience maps and the receptor/transporter density distributions. The statistical significance of the observed correlations was determined via permutation testing with 1,000 iterations, and p-values were derived from the null distribution. Multiple comparisons were corrected using the Bonferroni method.

## Results

### Higher emotional resilience among older adults

The demographic information, emotional ratings of movies, and behavioral measures of the current sample are summarized in **Table 1**. For emotional ratings, there was no significant group difference in valence rating before or after watching neutral movie (*p*s > 0.05), whereas older adults reported lower valence after watching the negative movie (*t* = 2.21, *p* = 0.029, *d* = 0.381). Regarding arousal ratings, older adults reported lower baseline arousal (*t* = 2.18, *p* = 0.031, *d* = 0.375), but no significant group differences in the neutral or negative movie conditions (*p*s > 0.05).

**Table 1.** Demographic information and studied variables.

|  | Younger Group<br>(N=68) | Older Group<br>(N=66) | *t*-value |
| --- | --- | --- | --- |
| Age(yrs) | 25.91 ± 4.31 | 71.02 ± 3.81 | - |
| TIV (cm<sup>3</sup>) | 1451.23 ± 204.26 | 1507.67 ± 192.36 | -1.64 |
| Head motion (mm) |  |  |  |
| *Neutral movie* | 0.13 ± 0.08 | 0.20 ± 0.08 | -5.35*** |
| *Negative movie* | 0.13 ± 0.06 | 0.21 ± 0.09 | -6.40*** |
| Standardized ERI | -0.58 ± 1.03 | 0.57 ± 0.53 | -7.93*** |
| Standardized CFI | 0.46 ± 0.73 | -0.41 ± 0.82 | 6.52*** |
| EV rating |  |  |  |
| *Baseline* | 6.10 ± 1.31 | 6.20 ± 1.70 | -0.36 |
| *Neutral movie* | 6.01 ± 1.31 | 5.88 ± 1.80 | 0.50 |
| *Negative movie* | 3.87 ± 1.49 | 3.24 ± 1.78 | 2.21* |
| EA rating |  |  |  |
| *Baseline* | 4.87 ± 1.53 | 4.23 ± 1.87 | 2.18* |
| *Neutral movie* | 4.69 ± 1.91 | 4.15 ± 1.96 | 1.61 |
| *Negative movie* | 6.21 ± 1.57 | 6.03 ± 1.77 | 0.61 |
| Amygdala<br>volume(cm<sup>3</sup>) |  |  |  |
| *Whole Amygdala* | 3.37 ± 0.32 | 3.03 ± 0.35 | 5.81*** |
| *BLA* | 1.69 ± 0.17 | 1.54 ± 0.18 | 5.06*** |
| *CMA* | 0.89 ± 0.08 | 0.79 ± 0.09 | 6.71*** |
| *SFA* | 0.78 ± 0.08 | 0.70 ± 0.09 | 5.87*** |

Note: TIV=total intracranial volume; ERI=emotional resilience index; CFI = cognitive function index; BLA= basolateral amygdala; CMA= centromedial amygdala; SFA= superficial amygdala; EV=emotional valence; EA=emotional arousal; \*: $p<0.05$ , \*\*: $p<0.01$ , \*\*\*: $p<0.001$ .

The younger group exhibited lower score in the emotional resilience index (ERI; *t* = −7.93, *p* < 0.001, *d* = −1.37) and higher score in the cognitive function index (CFI; *t* = −6.52, *p* < 0.001, *d* = 1.12) than the older group (**Fig.S3A**), indicating that younger adults showed lower emotional resilience, yet better cognitive performance compared with older adults. Group-specific correlation analyses showed a significant positive relationship between ERI and CFI in older adults (*r* = 0.25, *p* = 0.046), but not in younger adults (*r* = −0.10, *p* = 0.415, **Fig.S3B**). A follow-up moderation analysis revealed a trend-level effect of age on the relationship between ERI and CFI (β = 0.150, *p* = 0.081), suggesting that age may exert a modest influence on this association.

### Age and subfield specific effects

Whole-brain PLS analysis was conducted to identify age and subfield specific connectivity patterns for both negative and neutral movies. During the negative movie, first LV accounted for 63% of variance (*p* = 0.02) and revealed a pattern that was more activated among younger adults, similar across all subfields. Older adults recruited orthogonal and a different network, which was similarly engaged for all subfields (CIs overlapping). This network included anterior cingulate, bilateral inferior frontal gyrus, bilateral amygdala and hippocampus, bilateral thalamus, and bilateral putamen for younger adults. Regions such as right medial frontal gyrus, left inferior frontal gyrus, and left inferior temporal gyrus were engaged by older adults (**Fig.1A**). Second LV explained 26% variance of the data (*p* < 0.001) and included a network that was connected to BLA similarly for both age groups, although significantly more so among older adults (CI do not overlap). This network included left inferior frontal gyrus, bilateral middle frontal gyrus, bilateral insula, bilateral amygdala, bilateral parahippocampus, bilateral fusiform gyrus, and left superior temporal gyrus regions (**Fig.1B**).

**Figure. 1.**
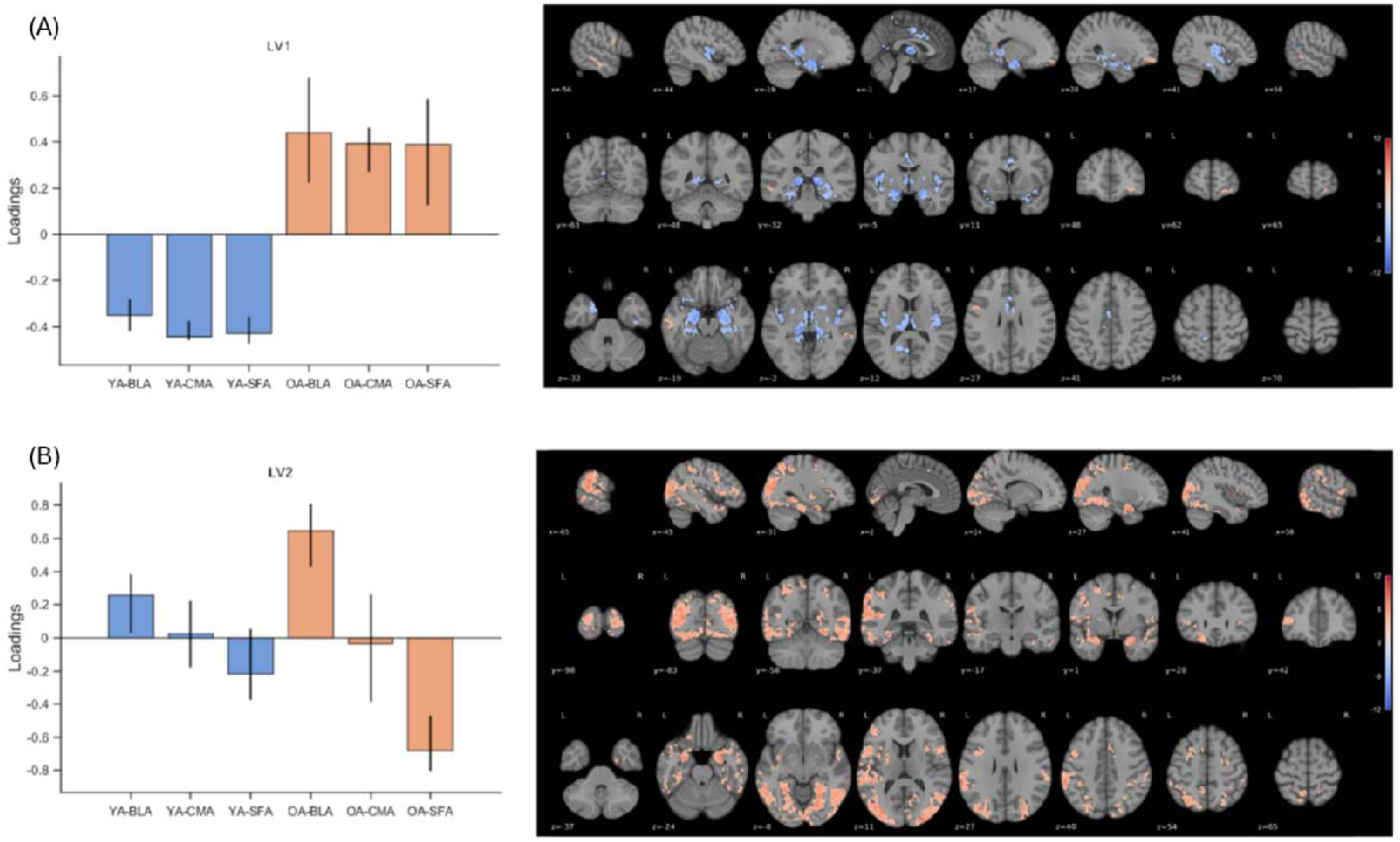
Functional networks associated with amygdala subfields across age groups during the negative movie. **(A)** LV1 represents the overlap across all amygdala subfields engaged in older adults. The salience map shows areas engaged by older adults (yellow) and younger adults (blue), similarly across all subfields. **(B)** LV2 shows similar activation of the BLA in both younger and older adults. The brain salience map indicates regions that were positively connected with the BLA in both age groups. The map was thresholded at bootstrap ratios (BSR) ≥ |3.3| (approximately p < 0.001), with a minimum cluster size of 400 mm³ (≈205 voxels at 1.25-mm resolution).

As for the neutral movie, first LV accounted for 68% variance of the data (*p* < 0.001) and engaged a network that was similarly engaged for all amygdala subfields among older adults. This network included bilateral anterior and posterior cingulate gyrus, bilateral inferior frontal gyrus, bilateral superior frontal gyrus, bilateral insula, bilateral inferior temporal gyrus, bilateral inferior parietal lobule, and bilateral precuneus. Younger adults, however, recruited an orthogonal pattern that included bilateral amygdala, bilateral hippocampus, bilateral thalamus, and right insula regions, which were similarly connected to all subfields of the amygdala (**Fig.S4A**). Second LV accounted for 23% of variance of the data (*p* = 0.007) and included right anterior cingulate, bilateral insula, bilateral superior temporal gyrus, bilateral thalamus, bilateral posterior cingulate gyrus, bilateral hippocampus, bilateral amygdala and putamen regions. These regions were connected to BLA similarly for both younger and older adults’ groups (**Fig.S4B**).

### Age impacts the association between amygdala subfields and behavioral markers

Next, we conducted brain-behavior PLS analyses to identify patterns of amygdala functional connectivity associated with ERI and CFI in younger and older adults. Specifically, three separate PLS analyses were performed using functional connectivity maps derived from each amygdala subregion (SFA, BLA, and CMA) as seed regions during the negative condition.

The analysis with BLA revealed one significant LV that accounted for 56% of the variance in the data (*p* = 0.005). The pattern reflected a distributed network covering the inferior and superior frontal gyrus, anterior cingulate cortex, insula, precuneus, and superior temporal gyrus. Increased connectivity between BLA and this network was positively correlated with higher ERI and CRI in older adults, and higher CRI and lower ERI in younger adults (**Fig.2**).

**Figure. 2.**
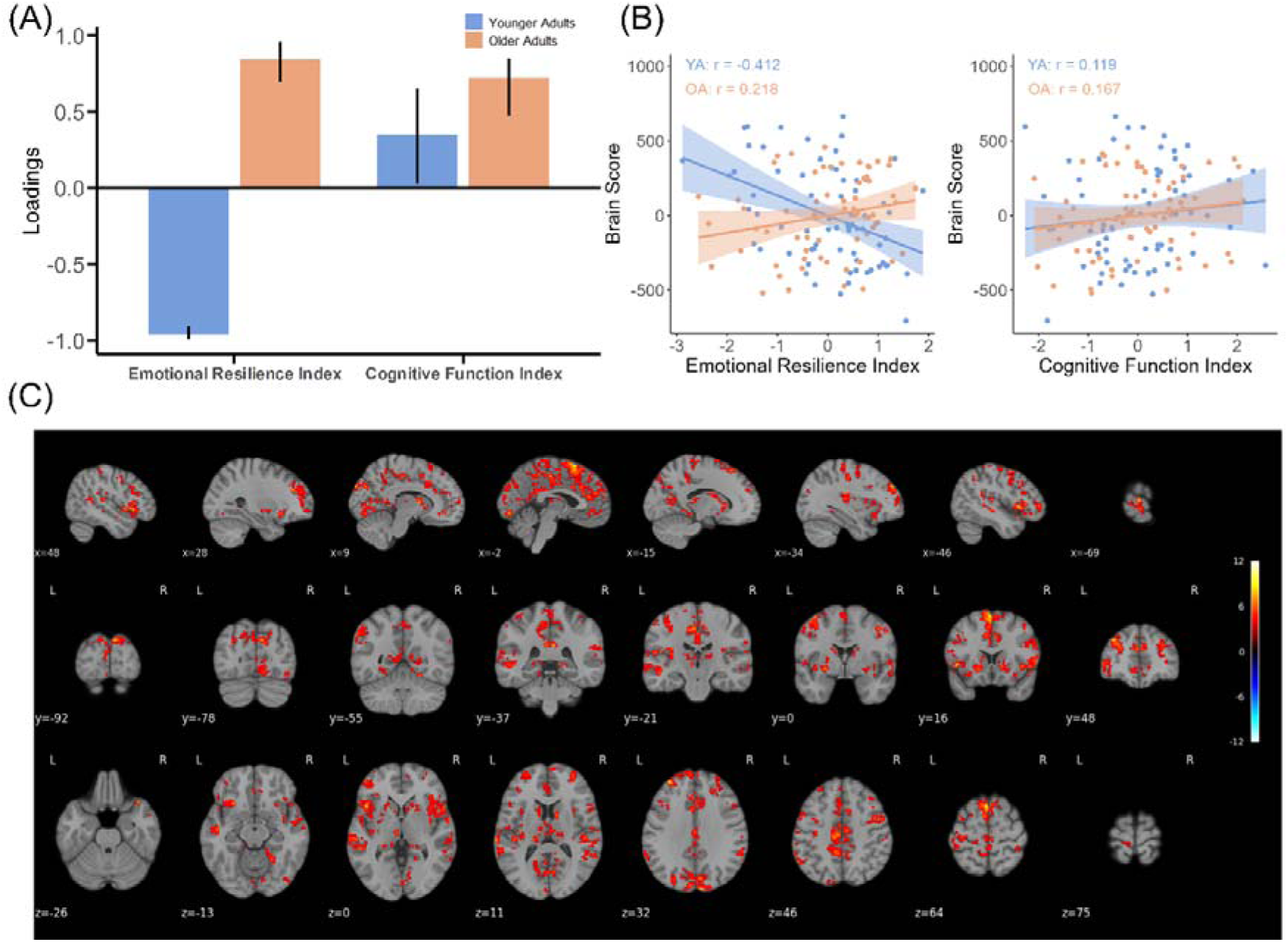
Brain–behavior associations for basolateral amygdala (BLA). **(A)** Behavioral loadings for each variable contributing to the significant latent variable (LV1). Error bars represent 95% confidence intervals derived from bootstrap resamples. Confidence intervals that do not include zero are interpreted as reliably contributing to the identified brain–behavior pattern. **(B)** Scatterplots show the correlations between brain scores and the emotional resilience index and cognitive function index in two age groups. **(C)** Salience map showing the BLA-related functional network associated with behavioral indices. The map was thresholded at bootstrap ratios (BSR) ≥ |3.3| (approximately p < 0.001) with a minimum cluster size of 400 mm³ (≈205 voxels at 1.25-mm resolution).

The analysis with the CMA revealed one significant LV that accounted for 60% of the variance in the data (*p* < 0.001). The identified brain pattern involved widespread regions spanning the medial prefrontal cortex, inferior parietal lobe, precuneus, posterior cingulate cortex, superior frontal gyrus, and middle temporal gyrus. Among younger adults, stronger FC between the CMA and these regions was linked to lower ERI, whereas no reliable association was found with CFI. Conversely, in older adults, an opposite pattern emerged with higher CMA network being associated with greater ERI and CFI (**Fig.3**).

**Figure. 3.**
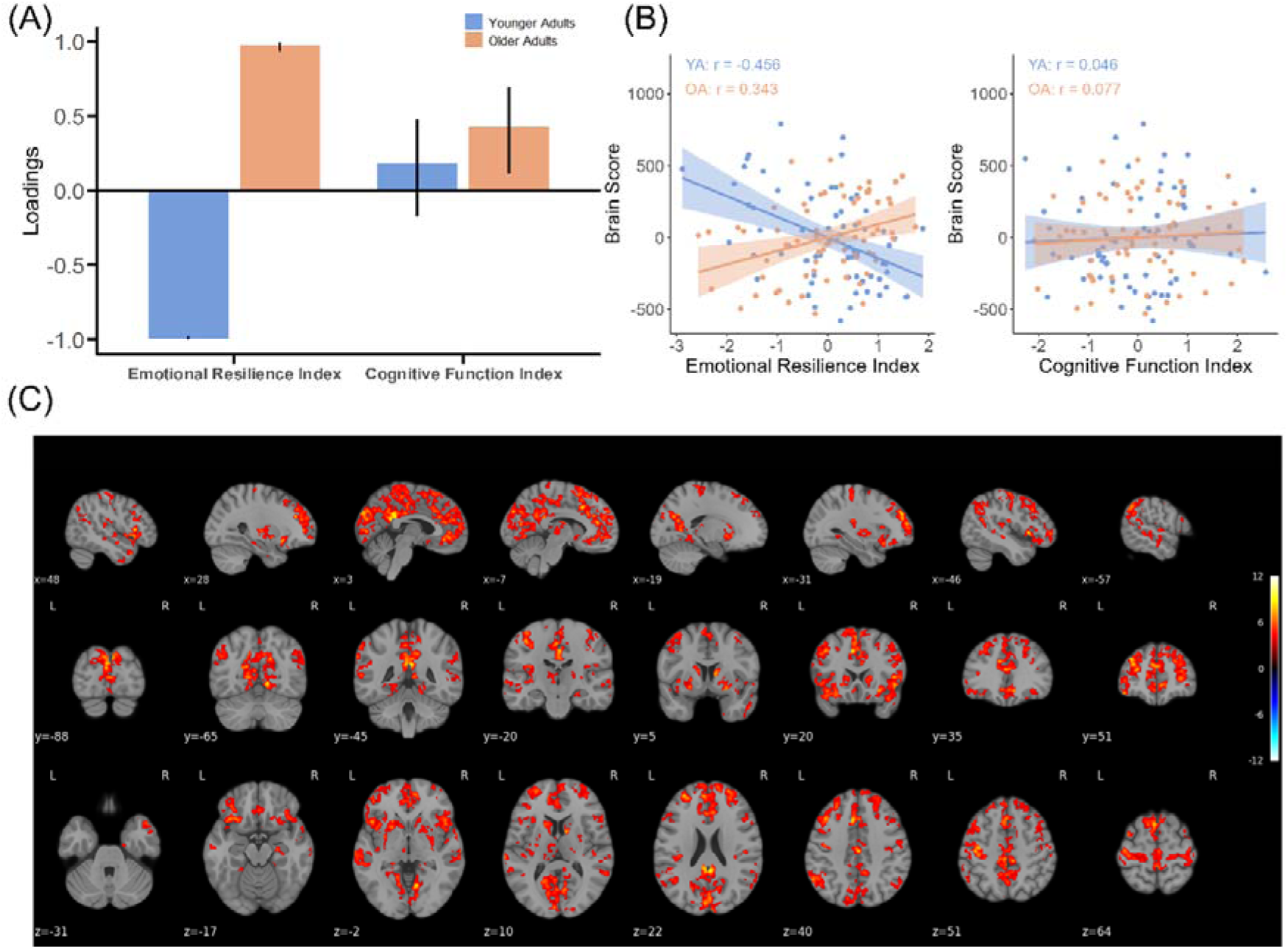
Brain–behavior associations for centromedial amygdala (CMA). **(A)** Behavioral loadings for each variable contributing to the significant latent variable (LV1). Error bars represent 95% confidence intervals derived from bootstrap resamples. Confidence intervals that do not include zero are interpreted as reliably contributing to the identified brain–behavior pattern. **(B)** Scatterplots show the correlations between brain scores and the emotional resilience index and cognitive function index in two age groups. **(C)** Salience map showing the CMA-related functional network associated with behavioral indices which was reliably activated by older adults. The map was thresholded at bootstrap ratios (BSR) ≥ |3.3| (approximately p < 0.001) with a minimum cluster size of 400 mm³ (≈205 voxels at 1.25-mm resolution).

The analysis with SFA resulted in only one significant LV which accounted for 55% of the variance of the data (*p* = 0.006). The salience map revealed a widespread pattern involving the anterior cingulate cortex, inferior and middle frontal gyrus, insula, precentral gyrus, superior temporal gyrus, and cuneus. Similar to the findings for the CMA, higher connectivity between the SFA and these brain regions was associated with lower CRI in younger adults, but with higher ERI and CFI in older adults (**Fig.4**). Additional analyses under the neutral condition identified no significant latent variables across the amygdala or its subregions, suggesting specificity of these findings to emotionally driven condition. Full results are presented in **Tab.S1**.

**Figure. 4.**
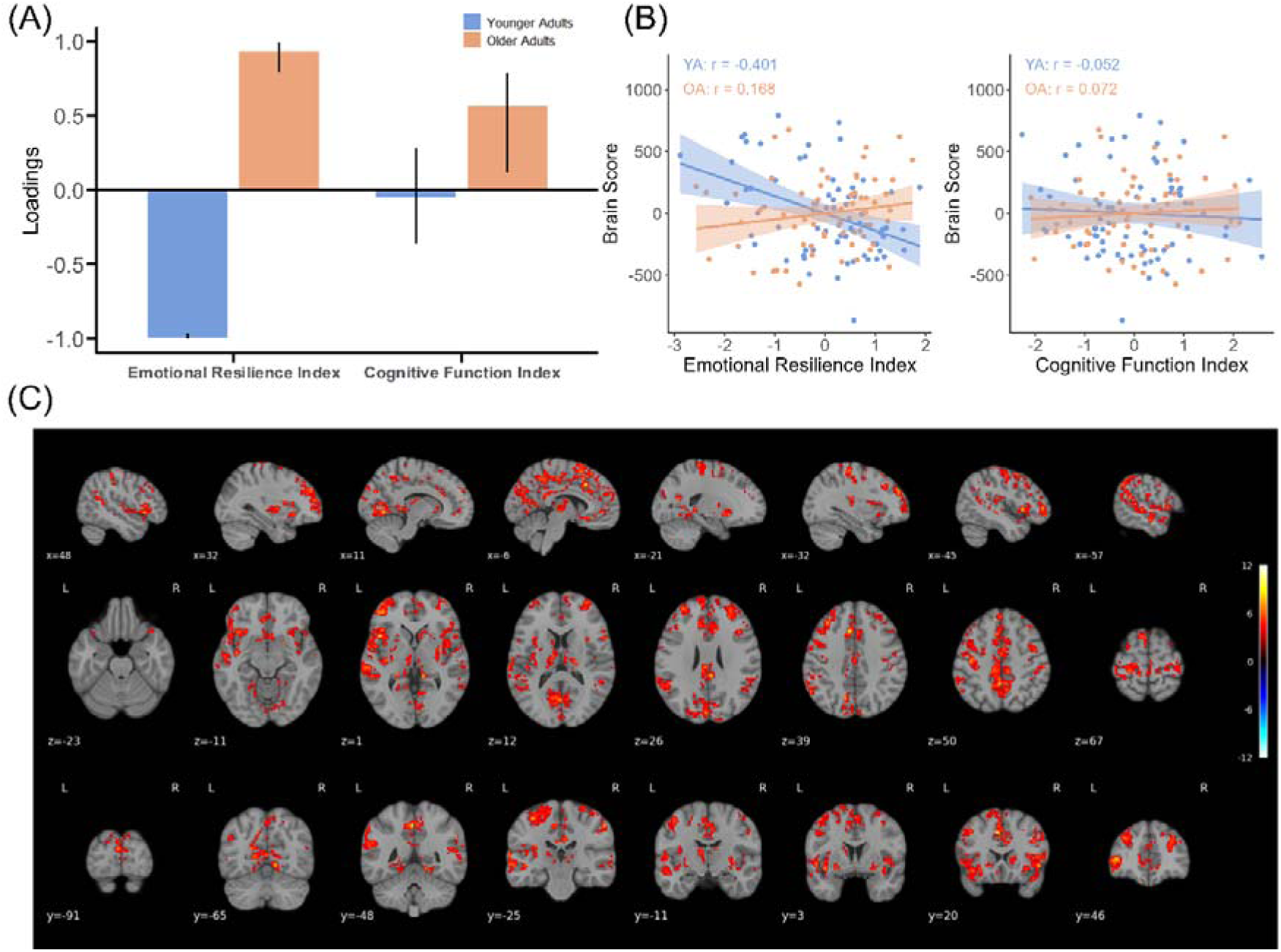
Brain–behavior associations for superficial amygdala (SFA). **(A)** Behavioral loadings for each variable contributing to the significant latent variable (LV1). Error bars represent 95% confidence intervals derived from bootstrap resamples. Confidence intervals that do not include zero are interpreted as reliably contributing to the identified brain–behavior pattern. **(B)** Scatterplots show the correlations between brain scores and the emotional resilience index and cognitive function index in two age groups. **(C)** Salience map showing the SFA-related functional network associated with behavioral indices which was activated by older adults. The map was thresholded at bootstrap ratios (BSR) ≥ |3.3| (approximately p < 0.001) with a minimum cluster size of 400 mm³ (≈205 voxels at 1.25-mm resolution).

A control analysis was conducted to examine the connectivity profile of the whole amygdala in relation to ERI and CFI, which yielded findings consistent with those observed in the subregional analyses and are presented in **Fig.S5**. Dice similarity coefficients between salience maps of the amygdala and its subregions ranged from 0.55 to 0.74, indicating a moderate-to-high spatial overlap with preserved regional specificity (**Fig.S6**).

Overall, our results indicate that the functional networks of the amygdala subregions associated with ERI and CFI largely overlap, although the strength and direction of these associations differ across age groups. Notably, these patterns emerged only during the negative movie condition. To further elucidate the network characteristics associated with ERI and CFI, additional analyses were performed to examine the overlapping and distinct connectivity profiles of the functional networks of each amygdala subregions.

### Overlap and distinct network correspondence of Amygdala subregion

To examine the overlapping between canonical functional networks and the networks connected to the amygdala subregions, network correspondence analysis was conducted. The results were present in **Fig.5**. Because the network labels differ across atlases, we adopted the abbreviations used in the toolbox to ensure consistency and reproducibility, rather than relying on broader large-scale network labels. The full network names corresponding to these abbreviations are provided in **Fig.S2**.

**Figure 5.**
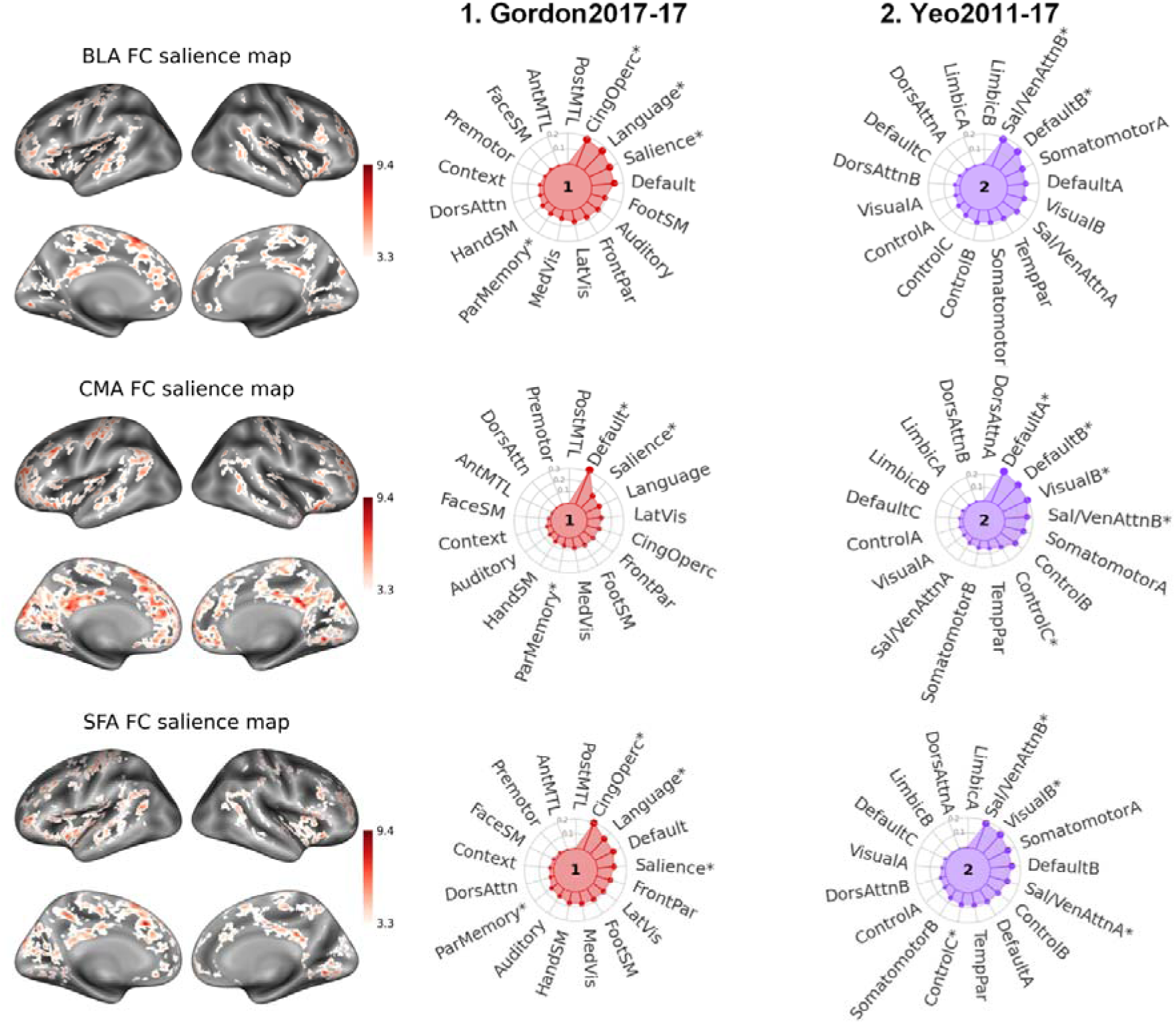
Network correspondence of amygdala subregion functional connectivity salience maps. Network radar plots display the Dice coefficients across canonical functional networks from two atlases, Gordon2017-17 and Yeo2011-17. Functional networks showing significant overlap with the salience map (spin test, FDR-corrected *p*□<□0.05) are indicated by asterisks.

The BLA salience map overlapped with CingOperc (Dice coefficient = 0.18, *p* = 0.022), Language (Dice coefficient = 0.17, *p* = 0.013), Salience (Dice coefficient = 0.14, *p* = 0.001), and ParMemory networks (Dice coefficient = 0.05, *p* = 0.048) in the Gordon2017-17 atlas, and with Sal/VenAttnB (Dice coefficient = 0.19, *p* = 0.002) and DefaultB (Dice coefficient = 0.17, *p* = 0.019) in the Yeo2011-17 atlas.

The salience map obtained from the PLS analysis focusing on the CMA revealed significant overlaps with the Default, Salience, and ParMemory networks in the Gordon2017-17 atlas, as well as with the DefaultA, DefaultB, VisualB, and ControlC networks in the Yeo2011-17 atlas.

For the SFA, the salience map showed significant overlap with the CingOperc (Dice coefficient = 0.20, *p* = 0.006), Language (Dice coefficient = 0.14, *p* = 0.038), Salience (Dice coefficient = 0.11, *p* = 0.001), and ParMemory networks (Dice coefficient = 0.06, *p* = 0.020) in the Gordon2017-17 atlas, as well as with the Salience/Ventral attention (Dice coefficient = 0.18, *p* = 0.002), Visual (Dice coefficient = 0.17, *p* = 0.003), Sal/VenAttnA (Dice coefficient = 0.12, *p* = 0.012) and ContralC networks in the Yeo2011-17 atlas.

### Functional decoding *of* amygdala subregion networks

Functional decoding revealed both shared and distinct functional profiles across the salience map of amygdala subregions (**Fig.6A**). The BLA was associated with inhibition, language comprehension, emotion regulation, and context. The CMA was primarily linked to emotion regulation, risk, mood, autobiographical memory. The SFA showed stronger associations with terms such as inhibition, monitoring, pain, and attention.

**Figure 6.**
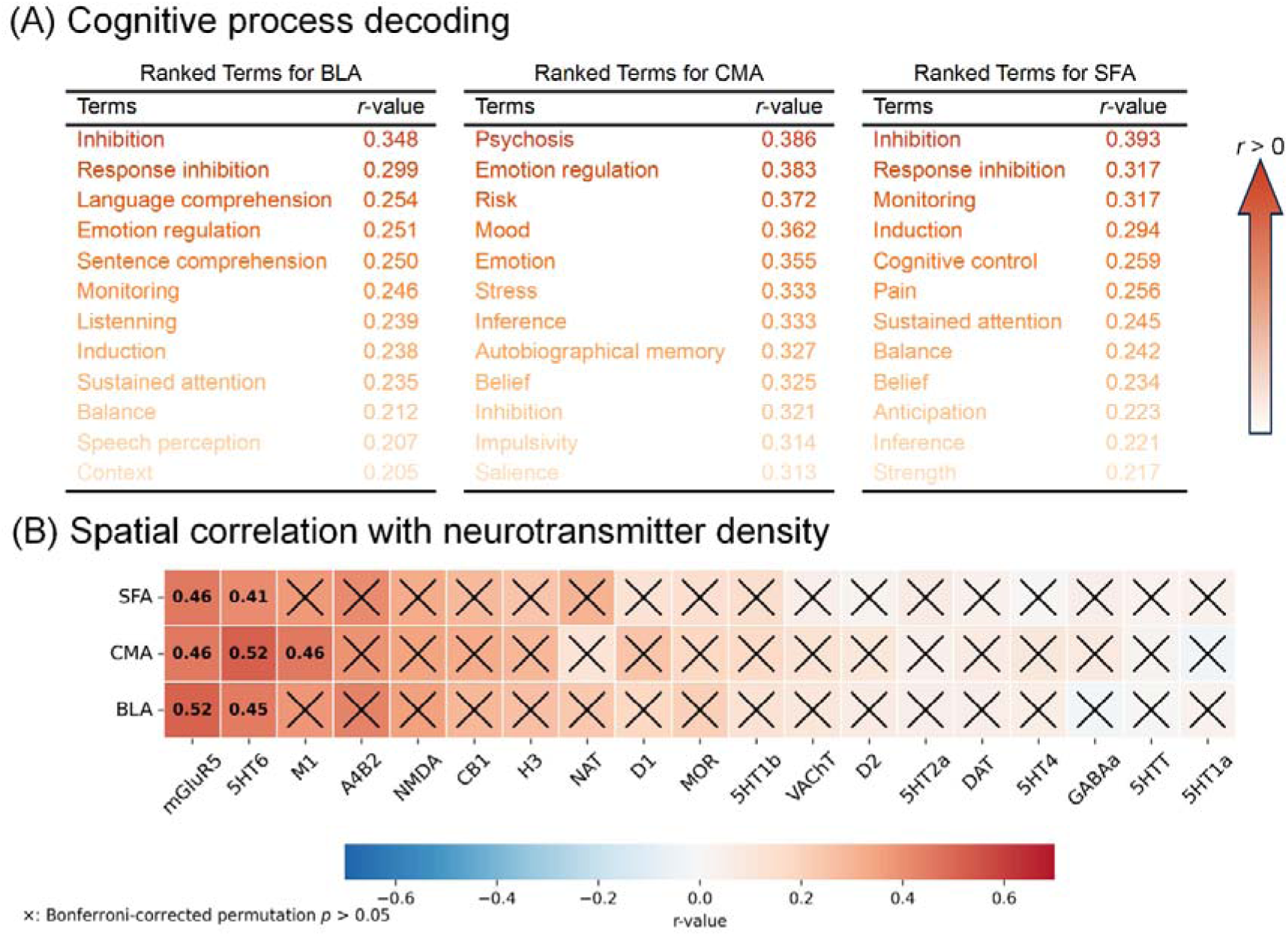
Cognitive process decoding of the salience maps and their spatial correspondence with neurotransmitter density distributions. **(A)** Cognitive process decoding of the salience maps derived from PLS analyses for the BLA, CMA, and SFA. Each column lists the top-ranked cognitive terms (based on r-values) showing the strongest associations with the identified connectivity patterns. **(B)** The heatmap shows the associations between the salience maps of each amygdala subregion and neurotransmitter receptor/transporter density maps.

### Neurotransmitter correspondence of amygdala subregion networks

We further examined the associations between the salience maps and neurotransmitter density maps to identify the underlying neurochemical mechanisms that may contribute to the observed brain–behavior relationships (**Fig.6B**). The networks of all three amygdala subregions were significantly correlated with spatial distribution of mGluR5 (BLA: *r* = 0.516, *p_corrected_* = 0.006; CMA: *r* = 0.463, *p_corrected_* = 0.006; SFA: *r* = 0.463, *p_corrected_* = 0.023) and 5HT6 (BLA: *r* = 0.452, *p_corrected_* = 0.006; CMA: *r* = 0.520, *p_corrected_* = 0.006; SFA: *r* = 0.411, *p_corrected_* = 0.006BLA: *r* = 0.452, *p_corrected_* = 0.006). Moreover, the salience map of CMA was significantly correlated with density of M1 (*r* = 0.461, *p_corrected_* = 0.006). These results suggest that the mGluR5 and 5HT6 might be potential candidates explaining the age-related differences in the amygdala network architecture and its association with behavioral phenotypes.

## Discussion

In this study, we combined ultra-high-field 7T fMRI with a naturalistic movie-watching paradigm to examine how emotional and cognitive functions relate to the functional profile of amygdala subregions’ connectivity in both younger and older adults. Using an ecologically and emotionally rich context, we examined age and subfields’ effects for the connectivity of amygdala subregions and investigated association between functional connectivity and behavioral correlates using a multivariate approach. Across all three subregions, BLA, CMA, and SFA, the analyses revealed distributed cortical and subcortical connectivity patterns that were systematically related to individual differences in emotional resilience and cognitive reserve indices, with clear age-dependent dissociations. The identified patterns mapped onto large-scale cortical systems—including the salience, control, and default mode network, highlighting the amygdala’s integrative role in bridging perceptual, regulatory, and self-referential processes that jointly sustain psychological well-being and cognitive functions.

We found strong overlap but also distinct functional profiles across all three amygdala subfields. The BLA and SFA exhibited overlapping connectivity profiles encompassing the anterior cingulate, insula, inferior and superior frontal gyri, superior temporal regions, precuneus, and angular gyrus—regions spanning across the salience, cingulo-opercular, control, language/ventral attention, and parietal memory networks, as identified through the network correspondence toolbox (Dosenbach et al., 2008; Gilmore et al., 2015; Menon & Uddin, 2010; Pessoa, 2017). Neurosynth decoding reinforced this interpretation by highlighting a consistent association with inhibition, emotion regulation, conflict monitoring, sustained attention, and broader cognitive control processes. Together, these functions describe a domain-general control system that supports adaptive emotion–cognition integration by linking amygdala signals, specifically from BLA and SFA, with salience detection and executive regulation. The contribution of the language–ventral attention network, which anchored part of this overlap, suggests an additional role for semantic appraisal and flexible attentional shifts in shaping context-appropriate emotional responses (Corbetta & Shulman, 2002; Fedorenko et al., 2011; Vossel et al., 2014).

These networks’ coupling with the amygdala are aligned with prior literature and likely enables efficient detection of emotionally relevant cues via insula–cingulate hubs (Alexandra Kredlow et al., 2022; Zhang et al., 2022), recruitment of prefrontal regions for top-down inhibition and monitoring (Gold et al., 2015; W.-Z. Liu et al., 2020), and coordination with parietal and temporal systems for contextual interpretation of affective information (Janak & Tye, 2015; Ueno et al., 2020). The cingulo-opercular network (CON), implicated in error monitoring, regulatory set maintenance, and sustained control (Dosenbach et al., 2007; Sadaghiani & D’Esposito, 2015), thus plays a key role in maintaining affective stability during prolonged emotional demands in the emotional movie condition. The inclusion of parietal memory regions, notably the precuneus and angular gyrus—indicates that BLA and SFA connectivity extends beyond rapid salience detection to incorporate emotionally guided retrieval and memory-based reappraisal, consistent with age-related reliance on posterior midline and parietal networks during emotional memory retrieval (Kensinger & Schacter, 2008; Ritchey et al., 2015; Spaniol & Grady, 2012).

In terms of association with emotional resilience scores, stronger BLA and SFA coupling with these distributed networks was associated with higher scores among older adults, suggesting that enhanced cortical integration between these subfields supports sustained regulatory be careful by engagement and emotional appraisal in late life. Such coactivation has been linked to the positivity affect, suggesting that increased engagement of memory and control networks supports selective regulation of emotionally meaningful material in older adults (Carstensen et al., 2011; Mather, 2016; Ziaei et al., 2016, 2017). In contrast, younger adults showed the opposite trend, stronger BLA/SFA connectivity predicted lower emotional resilience suggesting that extensive coupling between the amygdala and salience/attention systems may reflect heightened emotional reactivity or inefficient control (St. Jacques et al., 2010), particularly when affective networks operate in a more segregated and responsive mode. This pattern parallels clinical findings in PTSD and anxiety, where amygdala hyperconnectivity with the insula, cingulate, and VAN regions are associated with hypervigilance, intrusive re-experiencing, and affective dysregulation (MacDonald et al., 2025; Rabinak et al., 2011). In aging, however, similar configurations likely reflect functional reorganization, supporting compensatory engagement of attentional and control mechanisms to maintain emotional balance (Meunier et al., 2014; Morcom & Johnson, 2015).

The CMA exhibited more internally oriented connectivity pattern, linking the medial prefrontal, posterior cingulate, and inferior parietal cortices within the default mode and parietal memory networks, while maintaining connections with salience hubs in the anterior insula and dorsal anterior cingulate. Functional decoding associated this pattern with risk evaluation, mood regulation, emotion, stress processing, inhibition, inference, and autobiographical memory, suggesting that the CMA supports integration of visceral salience with introspective and evaluative functions (Barrett & Satpute, 2019; M.-S. Kong & Zweifel, 2021). In older adults, stronger CMA connectivity with DMN regions was associated with higher emotional and cognitive resilience, consistent with its role in stabilizing internal states and updating affective meaning through self-referential processing (Andrews-Hanna et al., 2014; Ronde et al., 2024; Spreng et al., 2010). Concurrent coupling with the salience network likely facilitates adaptive monitoring of internal arousal and dynamic regulatory control, while engagement of parietal memory nodes (angular gyrus, precuneus) enables access to autobiographical context for goal-consistent regulation (Gilmore et al., 2015; Ritchey & Cooper, 2020).

Together, these findings indicate that amygdala subfields–cortical network connectivity relates differentially to resilience across age. In older adults, broader integration of the BLA and SFA with salience, cingulo-opercular, and memory networks, and of the CMA with DMN–salience–memory systems, appears to support adaptive regulation, attentional stability, and meaning-based reappraisal (Sambuco, 2024; Seeley, 2019b; F. Zhou & Becker, 2025). In younger adults, by contrast, stronger coupling may reflect heightened affective sensitivity or less efficient inhibitory control, consistent with more modular network organization in early adulthood (Deery et al., 2023; Song et al., 2014). Thus, the partially distinct yet convergent roles of the BLA, SFA, and CMA suggest that the same patterns of amygdala connectivity may serve different functions across the lifespan, shifting from signals of heightened affective sensitivity in youth to markers of compensatory regulatory recruitment in older adults.

The overall amygdala-centered networks showed strong association with emotional resilience but only weak or not association with cognitive reserve across age groups. This is consistent with prior work identifying the amygdala as a central hub for emotional processing (Auer et al., 2025; Gothard, 2020). These results highlight the amygdala’s critical role in supporting emotional functions that help maintain psychological well-being. In contrast, for cognitive reserve, we only observed an association with the BLA-related network in younger adults, whereas older adults showed associations across all three networks of amygdala subnuclei. We speculate that this age-specific pattern may reflect functional dedifferentiation in aging (Bethlehem et al., 2020; Chan et al., 2014). That is, neural circuits traditionally dedicated to emotional processing may become increasingly recruited for cognitive processes in older adults. Alternatively, in line with compensatory accounts of aging, amygdala-based emotional circuits may be more heavily engaged during cognitive processes to offset declines in cognition-related brain systems (Morcom & Johnson, 2015). This interpretation aligns with our observation that emotional resilience and cognitive reserve were positively correlated only in the older adults group.

The neurochemical analyses further explained the amygdala network patterns by linking them to underlying neurotransmitter systems. All three amygdala subregion networks showed significant spatial coupling with mGluR5, a metabotropic glutamate receptor 5, and 5-HT6, serotonin neurotransmitter, receptor distributions, suggesting that glutamatergic and serotonergic modulation jointly shape the excitatory–inhibitory balance and plasticity of amygdala-centered circuits (Ligneul & Mainen, 2023; Y. Zhou & Danbolt, 2014). The mGluR5 is widely expressed in limbic and associative cortices and plays a critical role in synaptic plasticity, emotional learning, and adaptive regulation(Homayoun & Moghaddam, 2010; Neyman & Manahan-Vaughan, 2008). It has therefore been implicated in the pathophysiology of neurodegenerative diseases and affective disorders (DeLorenzo et al., 2015; Mecca et al., 2020). Previous studies have demonstrated mGluR5 dysregulation in patients with PTSD and major depressive disorder, especially within fronto-limbic regions (Esterlis et al., 2022). Moreover, increased mGluR5 availability has been observed in the dorsolateral and ventromedial prefrontal cortex of patients with major depressive disorder following eight weeks of vortioxetine, a serotonin reuptake inhibitor, treatment (B. Liu et al., 2025). Thus, the strong correspondence between the spatial distribution of mGluR5 and the PLS salience maps may suggest that behavior-related connectivity partly depend on the integrity of glutamatergic signaling that enables flexible updating of affective value and attentional priorities across changing contexts—processes that are essential for maintaining mental well-being when facing negative events. Serotonin modulates a range of human mental processes, including reward, affect, impulsivity, and cognitive flexibility (Cools et al., 2008; Schweighofer et al., 2008; Worbe et al., 2014). The observed association with 5-HT6 further highlights the serotonergic contribution to emotion–cognition interactions, aligning with pharmacological and imaging evidence linking 5-HT6 receptor density to symptoms of affective disorder and cognitive decline in aging (Chagraoui et al., 2025; Siwek et al., 2024; Wesołowska, 2010). Intriguingly, rodent studies show that 5-HT neurons influence amygdala circuitry via serotonin–glutamate cotransmission, indicating that the functional operations of the amygdala rely on the integrated actions of both neurotransmitter systems (Sengupta et al., 2017). Beyond these shared associations, the CMA network uniquely correlated with the spatial distribution of M1, muscarinic acetylcholine, and receptor density. Given that M1 receptors are predominant in cortical and hippocampal regions involved in memory and self-referential processing, this pattern may reflect a cholinergic pathway that supports integration between emotional valence and autobiographical memory (Manca et al., 2025; Sanda et al., 2024). Such modulation may facilitate the reappraisal of affective experiences and sustain positive mood regulation.

Overall, the involvement of mGluR5, 5-HT6, and M1 receptors provides a plausible neurochemical basis for the observed age-dependent reversal in amygdala–behavior coupling. Both glutamatergic and serotonergic signaling undergo gradual decline and receptor reconfiguration across the lifespan (Hernandez et al., 2018; Lorke et al., 2006; Mecca et al., 2021; Radhakrishnan et al., 2018), potentially altering the efficiency and balance of amygdala–network communication. Although we do not have age-specific data, these findings may suggest that age-related differences in functional connectivity patterns could partly be accounted by the shift in neurotransmitter density, although further research is needed to verify these results with study-specific measurements acquired for both age groups.

It should be noted that these associations were identified during the viewing of a negative movie rather than the neutral control. Negative emotional contexts are known to engage the amygdala more strongly and to elicit broader interactions with salience and control systems(Everaerd et al., 2015; Henckens et al., 2016; Kirk et al., 2022, 2023). Moreover, previous evidence has demonstrated that threat scenes induced increased activation in emotion-related brain regions, such as the anterior cingulate cortex and insula (Straube et al., 2010). Therefore, the present findings mainly reflect amygdala-centered circuits that are preferentially recruited under emotional challenge, illustrating how aging reshapes the coordination between the amygdala and the broader brain network during negative and stressful experiences. In contrast, the neutral movie condition did not evoke comparable effects, likely because its low emotional arousal produced weaker amygdala engagement and limited variability in affective responses (Cui et al., 2021). Without sufficient affective response, individual differences in emotional resilience and cognitive reserve may not translate into measurable changes in amygdala-related FC.

Several limitations should be acknowledged. First, the study included only neutral and negatively valanced movie clips and did not examine responses to positive emotional content. Although the inclusion of negative and neutral stimuli enabled the identification of affect-specific effects, the absence of positive conditions limits the generalizability of the findings across the full spectrum of emotional experiences (Greenberg et al., 2015). Future research should incorporate positive emotional contexts to determine whether the observed age-dependent patterns are specific to negative emotions or represent a more general reorganization of emotion regulation circuits. Second, the cross-sectional design precludes causal inference regarding age-related changes. Longitudinal data will be needed to clarify whether the observed connectivity differences reflect developmental trajectories or cohort effects. Finally, the neurotransmitter maps were derived from independent PET datasets for healthy adults (Dukart et al., 2021; Hansen et al., 2022). While spatial correspondence provides valuable insights into potential neurochemical substrates, these associations remain indirect and should be interpreted cautiously until verified with receptor-specific imaging in the same participants.

In conclusion, the findings indicate that emotional resilience is anchored in partially distinct amygdala subregional networks whose functional roles shift across adulthood. In younger adults, stronger amygdala–cortical coupling aligned with lower resilience, whereas in older adults the same configurations supported more adaptive emotional process. The combined network and neurochemical evidence point to coordinated engagement of salience, control, and memory systems further shaped by glutamatergic, serotonergic, and cholinergic signaling—as a key mechanism through which amygdala circuits help sustain emotional well-being in later life.

## Acknowledgment

We would like to thank radiographers and MR physicists at the 7T MR center at NTNU for their help during this project. We also would like to thank our participants for their time and effort during the experiment. We thank Stian Framvik, Avneesh Jain, Jae Hong, and Karina Tømmerdal for their help during data collection.

## Authorship contribution statement

**Shuer Ye:** Conceptualization, Methodology, Investigation, Formal analysis, Data curation, Visualization, Validation, Writing – original draft, Writing – review & editing. **Arjun Dave:** Data analyses, Investigation, Visualization, Writing – review & editing. **Xiaqing Lan:** Data analysis, Writing – review & editing. **Menno Witter:** Validation, Investigation, Writing – review & editing. **Alireza Salami:** Conceptualization, Investigation, Writing – review & editing. **Maryam Ziaei:** Conceptualization, Supervision, Project administration, Funding acquisition, Writing – original draft, Writing – review & editing.

## Ethics and Consent to Participate

Participants provided written informed consent. The research complied with the Declaration of Helsinki and had ethical approval from the regional committees for medical and health research ethics (REK-Midt, approval #390390).

## References

Abdi, H., & Williams, L. J. (2010). Principal component analysis. WIREs Computational Statistics, 2(4), 433–459. 10.1002/wics.101

Aghamohammadi-Sereshki, A., Hrybouski, S., Travis, S., Huang, Y., Olsen, F., Carter, R., Camicioli, R., & Malykhin, N. V. (2019). Amygdala subnuclei and healthy cognitive aging. Human Brain Mapping, 40(1), 34–52. 10.1002/hbm.24353

Alarcón, G., Cservenka, A., Rudolph, M. D., Fair, D. A., & Nagel, B. J. (2015). Developmental sex differences in resting state functional connectivity of amygdala sub-regions. NeuroImage, 115, 235–244. 10.1016/j.neuroimage.2015.04.013

Alexandra Kredlow, M., Fenster, R. J., Laurent, E. S., Ressler, K. J., & Phelps, E. A. (2022). Prefrontal cortex, amygdala, and threat processing: Implications for PTSD. Neuropsychopharmacology, 47(1), 247–259. 10.1038/s41386-021-01155-7

Allen, H. N., Bobnar, H. J., & Kolber, B. J. (2021). Left and right hemispheric lateralization of the amygdala in pain. Progress in Neurobiology, 196, 101891. 10.1016/j.pneurobio.2020.101891

Andrews-Hanna, J. R., Smallwood, J., & Spreng, R. N. (2014). The default network and self-generated thought: Component processes, dynamic control, and clinical relevance. Annals of the New York Academy of Sciences, 1316(1), 29–52. 10.1111/nyas.12360

Auer, H., Cabalo, D. G., Rodríguez-Cruces, R., Benkarim, O., Paquola, C., DeKraker, J., Wang, Y., Valk, S. L., Bernhardt, B. C., & Royer, J. (2025). From histology to macroscale function in the human amygdala. eLife, 13, RP101950. 10.7554/eLife.101950

Baez-Lugo, S., Deza-Araujo, Y. I., Maradan, C., Collette, F., Lutz, A., Marchant, N. L., Chételat, G., Vuilleumier, P., & Klimecki, O. (2023). Exposure to negative socio-emotional events induces sustained alteration of resting-state brain networks in older adults. Nature Aging, 3(1), 105–120. 10.1038/s43587-022-00341-6

Baron-Cohen, S., Wheelwright, S., Hill, J., Raste, Y., & Plumb, I. (2001). The “Reading the Mind in the Eyes” Test Revised Version: A Study with Normal Adults, and Adults with Asperger Syndrome or High-functioning Autism. Journal of Child Psychology and Psychiatry, 42(2), 241–251. 10.1111/1469-7610.00715

Barrett, L. F., & Satpute, A. B. (2019). Historical pitfalls and new directions in the neuroscience of emotion. Neuroscience Letters, 693, 9–18. 10.1016/j.neulet.2017.07.045

Bethlehem, R. A. I., Paquola, C., Seidlitz, J., Ronan, L., Bernhardt, B., Consortium, C.-C., & Tsvetanov, K. A. (2020). Dispersion of functional gradients across the adult lifespan. NeuroImage, 222, 117299. 10.1016/j.neuroimage.2020.117299

Bjelland, I., Dahl, A. A., Haug, T. T., & Neckelmann, D. (2002). The validity of the Hospital Anxiety and Depression Scale: An updated literature review. Journal of Psychosomatic Research, 52(2), 69–77. 10.1016/S0022-3999(01)00296-3

Buhr, K., & Dugas, M. J. (2002). The intolerance of uncertainty scale: Psychometric properties of the English version. Behaviour Research and Therapy, 40(8), 931–945. 10.1016/S0005-7967(01)00092-4

Bzdok, D., Laird, A. R., Zilles, K., Fox, P. T., & Eickhoff, S. B. (2013). An investigation of the structural, connectional, and functional subspecialization in the human amygdala. Human Brain Mapping, 34(12), 3247–3266. 10.1002/hbm.22138

Carstensen, L. L., Turan, B., Scheibe, S., Ram, N., Ersner-Hershfield, H., Samanez-Larkin, G. R., Brooks, K. P., & Nesselroade, J. R. (2011). Emotional experience improves with age: Evidence based on over 10 years of experience sampling. Psychology and Aging, 26(1), 21–33. 10.1037/a0021285

Chagraoui, A., Thibaut, F., & De Deurwaerdère, P. (2025). 5-HT6 receptors: Contemporary views on their neurobiological and pharmacological relevance in neuropsychiatric disorders. Dialogues in Clinical Neuroscience, 27(1), 112–128. 10.1080/19585969.2025.2502028

Chan, M. Y., Park, D. C., Savalia, N. K., Petersen, S. E., & Wig, G. S. (2014). Decreased segregation of brain systems across the healthy adult lifespan. Proceedings of the National Academy of Sciences, 111(46), E4997–E5006. 10.1073/pnas.1415122111

Cohen, S., Kamarck, T., & Mermelstein, R. (1983). A Global Measure of Perceived Stress. Journal of Health and Social Behavior, 24(4), 385–396. 10.2307/2136404

Connor, K. M., & Davidson, J. R. T. (2003). Development of a new resilience scale: The Connor-Davidson Resilience Scale (CD-RISC. Depress. Anxiety, 18, 76–82.

Cools, R., Roberts, A. C., & Robbins, T. W. (2008). Serotoninergic regulation of emotional and behavioural control processes. Trends in Cognitive Sciences, 12(1), 31–40. 10.1016/j.tics.2007.10.011

Corbetta, M., & Shulman, G. L. (2002). Control of goal-directed and stimulus-driven attention in the brain. Nature Reviews Neuroscience, 3(3), 201–215. 10.1038/nrn755

Cui, L., Guo, B., Zhao, D., Li, J., Luo, Y., & Meng, M. (2021). Amygdala-based Functional Network Reveals Dissociated Neural Correlates of Consensual and Idiosyncratic Emotional Movie Experiences. Neuroscience Bulletin, 37(5), 729–734. 10.1007/s12264-021-00666-z

Dall’Oglio, A., Dutra, A. C. L., Moreira, J. E., & Rasia-Filho, A. A. (2015). The human medial amygdala: Structure, diversity, and complexity of dendritic spines. Journal of Anatomy, 227(4), 440–459. 10.1111/joa.12358

Damoiseaux, J. S. (2017). Effects of aging on functional and structural brain connectivity. NeuroImage, 160, 32–40. 10.1016/j.neuroimage.2017.01.077

Dave, A., Ye, S., Bätz, L. R., Lan, X., Jacobs, H. I. L., & Ziaei, M. (2025). Age-Related Increase in Locus Ceruleus Activity and Connectivity with the Prefrontal Cortex during Ambiguity Processing. Journal of Neuroscience, 45(32). 10.1523/JNEUROSCI.2059-24.2025

Davis, M. H. (1983). Measuring individual differences in empathy: Evidence for a multidimensional approach. Journal of Personality and Social Psychology, 44(1), 113–126. 10.1037/0022-3514.44.1.113

Deery, H. A., Di Paolo, R., Moran, C., Egan, G. F., & Jamadar, S. D. (2023). The older adult brain is less modular, more integrated, and less efficient at rest: A systematic review of large-scale resting-state functional brain networks in aging. Psychophysiology, 60(1), e14159. 10.1111/psyp.14159

DeLorenzo, C., Sovago, J., Gardus, J., Xu, J., Yang, J., Behrje, R., Kumar, J. S. D., Devanand, D. P., Pelton, G. H., Mathis, C. A., Mason, N. S., Gomez-Mancilla, B., Aizenstein, H., Mann, J. J., & Parsey, R. V. (2015). Characterization of brain mGluR5 binding in a pilot study of late-life major depressive disorder using positron emission tomography and [11C]ABP688. Translational Psychiatry, 5(12), e693–e693. 10.1038/tp.2015.189

Dolcos, F., Katsumi, Y., Moore, M., Berggren, N., de Gelder, B., Derakshan, N., Hamm, A. O., Koster, E. H. W., Ladouceur, C. D., Okon-Singer, H., Pegna, A. J., Richter, T., Schweizer, S., Van den Stock, J., Ventura-Bort, C., Weymar, M., & Dolcos, S. (2020). Neural correlates of emotion-attention interactions: From perception, learning, and memory to social cognition, individual differences, and training interventions. Neuroscience & Biobehavioral Reviews, 108, 559–601. 10.1016/j.neubiorev.2019.08.017

Dosenbach, N. U. F., Fair, D. A., Cohen, A. L., Schlaggar, B. L., & Petersen, S. E. (2008). A dual-networks architecture of top-down control. Trends in Cognitive Sciences, 12(3), 99–105. 10.1016/j.tics.2008.01.001

Dosenbach, N. U. F., Fair, D. A., Miezin, F. M., Cohen, A. L., Wenger, K. K., Dosenbach, R. A. T., Fox, M. D., Snyder, A. Z., Vincent, J. L., Raichle, M. E., Schlaggar, B. L., & Petersen, S. E. (2007). Distinct brain networks for adaptive and stable task control in humans. Proceedings of the National Academy of Sciences, 104(26), 11073–11078. 10.1073/pnas.0704320104

Dukart, J., Holiga, S., Rullmann, M., Lanzenberger, R., Hawkins, P. C. T., Mehta, M. A., Hesse, S., Barthel, H., Sabri, O., Jech, R., & Eickhoff, S. B. (2021). JuSpace: A tool for spatial correlation analyses of magnetic resonance imaging data with nuclear imaging derived neurotransmitter maps. Human Brain Mapping, 42(3), 555–566. 10.1002/hbm.25244

Efron, B., & Tibshirani, R. (1985). The Bootstrap Method for Assessing Statistical Accuracy. Behaviormetrika, 12(17), 1–35. 10.2333/bhmk.12.17_1

Eickhoff, S. B., Milham, M., & Vanderwal, T. (2020). Towards clinical applications of movie fMRI. NeuroImage, 217, 116860. 10.1016/j.neuroimage.2020.116860

Esteban, O., Markiewicz, C. J., Blair, R. W., Moodie, C. A., Isik, A. I., Erramuzpe, A., Kent, J. D., Goncalves, M., DuPre, E., Snyder, M., Oya, H., Ghosh, S. S., Wright, J., Durnez, J., Poldrack, R. A., & Gorgolewski, K. J. (2019). fMRIPrep: A robust preprocessing pipeline for functional MRI. Nature Methods, 16(1), 111–116. 10.1038/s41592-018-0235-4

Esterlis, I., DeBonee, S., Cool, R., Holmes, S., Baldassari, S. R., Maruff, P., Pietrzak, R. H., & Davis, M. T. (2022). Differential Role of mGluR5 in Cognitive Processes in Posttraumatic Stress Disorder and Major Depression. Chronic Stress, 6, 24705470221105804. 10.1177/24705470221105804

Everaerd, D., Klumpers, F., van Wingen, G., Tendolkar, I., & Fernández, G. (2015). Association between neuroticism and amygdala responsivity emerges under stressful conditions. NeuroImage, 112, 218–224. 10.1016/j.neuroimage.2015.03.014

Fedorenko, E., Behr, M. K., & Kanwisher, N. (2011). Functional specificity for high-level linguistic processing in the human brain. Proceedings of the National Academy of Sciences, 108(39), 16428–16433. 10.1073/pnas.1112937108

Finn, E. S. (2021). Is it time to put rest to rest? Trends in Cognitive Sciences, 25(12), 1021–1032. 10.1016/j.tics.2021.09.005

Fjell, A. M., McEvoy, L., Holland, D., Dale, A. M., Walhovd, K. B., & Initiative, for the A. D. N. (2013). Brain Changes in Older Adults at Very Low Risk for Alzheimer’s Disease. Journal of Neuroscience, 33(19), 8237–8242. 10.1523/JNEUROSCI.5506-12.2013

Friston, K. J., Holmes, A. P., Worsley, K. J., Poline, J.-P., Frith, C. D., & Frackowiak, R. S. J. (1994). Statistical parametric maps in functional imaging: A general linear approach. Human Brain Mapping, 2(4), 189–210. 10.1002/hbm.460020402

Gaser, C., Dahnke, R., Thompson, P. M., Kurth, F., Luders, E., & the Alzheimer’s Disease Neuroimaging Initiative. (2024). CAT: A computational anatomy toolbox for the analysis of structural MRI data. GigaScience, 13, giae049. 10.1093/gigascience/giae049

Gilmore, A. W., Nelson, S. M., & McDermott, K. B. (2015). A parietal memory network revealed by multiple MRI methods. Trends in Cognitive Sciences, 19(9), 534–543. 10.1016/j.tics.2015.07.004

Gold, A. L., Morey, R. A., & McCarthy, G. (2015). Amygdala–Prefrontal Cortex Functional Connectivity During Threat-Induced Anxiety and Goal Distraction. Biological Psychiatry, 77(4), 394–403. 10.1016/j.biopsych.2014.03.030

Gordon, E. M., Laumann, T. O., Gilmore, A. W., Newbold, D. J., Greene, D. J., Berg, J. J., Ortega, M., Hoyt-Drazen, C., Gratton, C., Sun, H., Hampton, J. M., Coalson, R. S., Nguyen, A. L., McDermott, K. B., Shimony, J. S., Snyder, A. Z., Schlaggar, B. L., Petersen, S. E., Nelson, S. M., & Dosenbach, N. U. F. (2017). Precision Functional Mapping of Individual Human Brains. Neuron, 95(4), 791–807.e7. 10.1016/j.neuron.2017.07.011

Gothard, K. M. (2020). Multidimensional processing in the amygdala. Nature Reviews Neuroscience, 21(10), 565–575. 10.1038/s41583-020-0350-y

Gratz, K. L., & Roemer, L. (2004). Multidimensional Assessment of Emotion Regulation and Dysregulation: Development, Factor Structure, and Initial Validation of the Difficulties in Emotion Regulation Scale. Journal of Psychopathology and Behavioral Assessment, 26(1), 41–54. 10.1023/B:JOBA.0000007455.08539.94

Greenberg, T., Carlson, J. M., Rubin, D., Cha, J., & Mujica-Parodi, L. (2015). Anticipation of high arousal aversive and positive movie clips engages common and distinct neural substrates. Social Cognitive and Affective Neuroscience, 10(4), 605–611. 10.1093/scan/nsu091

Gunning, M. D., Denison, F. C., Stockley, C. J., Ho, S. P., Sandhu, H. K., & Reynolds, R. M. (2010). Assessing maternal anxiety in pregnancy with the State□Trait Anxiety Inventory (STAI): Issues of validity, location and participation. Journal of Reproductive and Infant Psychology, 28(3), 266–273. 10.1080/02646830903487300

Hansen, J. Y., Cauzzo, S., Singh, K., García-Gomar, M. G., Shine, J. M., Bianciardi, M., & Misic, B. (2024). Integrating brainstem and cortical functional architectures. Nature Neuroscience, 27(12), 2500–2511. 10.1038/s41593-024-01787-0

Hansen, J. Y., Shafiei, G., Markello, R. D., Smart, K., Cox, S. M. L., Nørgaard, M., Beliveau, V., Wu, Y., Gallezot, J.-D., Aumont, É., Servaes, S., Scala, S. G., DuBois, J. M., Wainstein, G., Bezgin, G., Funck, T., Schmitz, T. W., Spreng, R. N., Galovic, M., … Misic, B. (2022). Mapping neurotransmitter systems to the structural and functional organization of the human neocortex. Nature Neuroscience, 25(11), 1569–1581. 10.1038/s41593-022-01186-3

Heimer, L., & Van Hoesen, G. W. (2006). The limbic lobe and its output channels: Implications for emotional functions and adaptive behavior. Neuroscience & Biobehavioral Reviews, 30(2), 126–147. 10.1016/j.neubiorev.2005.06.006

Henckens, M. J. A. G., Klumpers, F., Everaerd, D., Kooijman, S. C., van Wingen, G. A., & Fernández, G. (2016). Interindividual differences in stress sensitivity: Basal and stress-induced cortisol levels differentially predict neural vigilance processing under stress. Social Cognitive and Affective Neuroscience, 11(4), 663–673. 10.1093/scan/nsv149

Hernandez, C. M., McQuail, J. A., Schwabe, M. R., Burke, S. N., Setlow, B., & Bizon, J. L. (2018). Age-Related Declines in Prefrontal Cortical Expression of Metabotropic Glutamate Receptors that Support Working Memory. eNeuro, 5(3). 10.1523/ENEURO.0164-18.2018

Homayoun, H., & Moghaddam, B. (2010). Group 5 metabotropic glutamate receptors: Role in modulating cortical activity and relevance to cognition. European Journal of Pharmacology, 639(1), 33–39. 10.1016/j.ejphar.2009.12.042

Isaacowitz, D. M., Wadlinger, H. A., Goren, D., & Wilson, H. R. (2006). Is there an age-related positivity effect in visual attention? A comparison of two methodologies. Emotion, 6(3), 511–516. 10.1037/1528-3542.6.3.511

Ivnik, R. J., Malec, J. F., Smith, G. E., Tangalos, E. G., & Petersen, R. C. (1996). Neuropsychological tests’ norms above age 55: COWAT, BNT, MAE token, WRAT-R reading, AMNART, STROOP, TMT, and JLO. The Clinical Neuropsychologist, 10(3), 262–278. 10.1080/13854049608406689

Jacob, Y., Morris, L. S., Verma, G., Rutter, S. B., Balchandani, P., & Murrough, J. W. (2022). Altered hippocampus and amygdala subregion connectome hierarchy in major depressive disorder. Translational Psychiatry, 12(1), 209. 10.1038/s41398-022-01976-0

Janak, P. H., & Tye, K. M. (2015). From circuits to behaviour in the amygdala. Nature, 517(7534), 284–292. 10.1038/nature14188

Jensen, A. R., & Rohwer, W. D. (1966). The stroop color-word test: A review. Acta Psychologica, 25, 36–93. 10.1016/0001-6918(66)90004-7

Kensinger, E. A., & Schacter, D. L. (2008). Neural Processes Supporting Young and Older Adults’ Emotional Memories. Journal of Cognitive Neuroscience, 20(7), 1161–1173. 10.1162/jocn.2008.20080

Kirk, P. A., Holmes, A. J., & Robinson, O. J. (2023). Anxiety Shapes Amygdala-Prefrontal Dynamics During Movie Watching. Biological Psychiatry Global Open Science, 3(3), 409–417. 10.1016/j.bpsgos.2022.03.009

Kirk, P. A., Robinson, O. J., & Skipper, J. I. (2022). Anxiety and amygdala connectivity during movie-watching. Neuropsychologia, 169, 108194. 10.1016/j.neuropsychologia.2022.108194

Kirstein, C. F., Güntürkün, O., & Ocklenburg, S. (2023). Ultra-high field imaging of the amygdala – A narrative review. Neuroscience & Biobehavioral Reviews, 152, 105245. 10.1016/j.neubiorev.2023.105245

Kong, M.-S., & Zweifel, L. S. (2021). Central amygdala circuits in valence and salience processing. Behavioural Brain Research, 410, 113355. 10.1016/j.bbr.2021.113355

Kong, R., Spreng, R. N., Xue, A., Betzel, R. F., Cohen, J. R., Damoiseaux, J. S., De Brigard, F., Eickhoff, S. B., Fornito, A., Gratton, C., Gordon, E. M., Holmes, A. J., Laird, A. R., Larson-Prior, L., Nickerson, L. D., Pinho, A. L., Razi, A., Sadaghiani, S., Shine, J. M., … Uddin, L. Q. (2025). A network correspondence toolbox for quantitative evaluation of novel neuroimaging results. Nature Communications, 16(1), 2930. 10.1038/s41467-025-58176-9

Krishnan, A., Williams, L. J., McIntosh, A. R., & Abdi, H. (2011). Partial Least Squares (PLS) methods for neuroimaging: A tutorial and review. NeuroImage, 56(2), 455–475. 10.1016/j.neuroimage.2010.07.034

Kurth, F., Cherbuin, N., & Luders, E. (2019). Age but no sex effects on subareas of the amygdala. Human Brain Mapping, 40(6), 1697–1704. 10.1002/hbm.24481

Kwon, H., Ha, M., Choi, S., Park, S., Jang, M., Kim, M., & Kwon, J. S. (2024). Resting-state functional connectivity of amygdala subregions across different symptom subtypes of obsessive–compulsive disorder patients. NeuroImage: Clinical, 43, 103644. 10.1016/j.nicl.2024.103644

Ligneul, R., & Mainen, Z. F. (2023). Serotonin. Current Biology, 33(23), R1216–R1221. 10.1016/j.cub.2023.09.068

Liu, B., Deng, A., Dong, C., Chen, W., Zhang, Q., Zhou, L., He, F., Xiang, X., Ou, W., Ma, M., Liu, J., Wang, X., Ju, Y., Wang, Y., Huang, H., Ma, X., & Zhang, Y. (2025). Enhanced mGluR5 availability marks the antidepressant efficacy in major depressive disorder: An [18F]FPEB PET study. Nature Mental Health, 3(3), 298–305. 10.1038/s44220-025-00386-7

Liu, W.-Z., Zhang, W.-H., Zheng, Z.-H., Zou, J.-X., Liu, X.-X., Huang, S.-H., You, W.-J., He, Y., Zhang, J.-Y., Wang, X.-D., & Pan, B.-X. (2020). Identification of a prefrontal cortex-to-amygdala pathway for chronic stress-induced anxiety. Nature Communications, 11(1), 2221. 10.1038/s41467-020-15920-7

Lorke, D. E., Lu, G., Cho, E., & Yew, D. T. (2006). Serotonin 5-HT2A and 5-HT6 receptors in the prefrontal cortex of Alzheimer and normal aging patients. BMC Neuroscience, 7(1), 36. 10.1186/1471-2202-7-36

Lovibond, P. F., & Lovibond, S. H. (1995). The structure of negative emotional states: Comparison of the Depression Anxiety Stress Scales (DASS) with the Beck Depression and Anxiety Inventories. Behaviour Research and Therapy, 33(3), 335–343. 10.1016/0005-7967(94)00075-U

Luppi, A. I., Golkowski, D., Ranft, A., Ilg, R., Jordan, D., Bzdok, D., Owen, A. M., Naci, L., Stamatakis, E. A., Amico, E., & Misic, B. (2025). General anaesthesia decreases the uniqueness of brain functional connectivity across individuals and species. Nature Human Behaviour, 9(5), 987–1004. 10.1038/s41562-025-02121-9

MacDonald, S. E., Becker, C. R., & MacNamara, A. (2025). Amygdala-insula response to neutral stimuli and the prospective prediction of anxiety sensitivity. Progress in Neuro-Psychopharmacology and Biological Psychiatry, 139, 111384. 10.1016/j.pnpbp.2025.111384

Manca, R., De Marco, M., Soininen, H., Ruffini, L., & Venneri, A. (2025). Changes in neurotransmitter-related functional connectivity along the Alzheimer’s disease continuum. Brain Communications, 7(1), fcaf008. 10.1093/braincomms/fcaf008

Mather, M. (2016). The Affective Neuroscience of Aging. Annual Review of Psychology, 67(Volume 67, 2016), 213–238. 10.1146/annurev-psych-122414-033540

McIntosh, A. R., Bookstein, F. L., Haxby, J. V., & Grady, C. L. (1996). Spatial Pattern Analysis of Functional Brain Images Using Partial Least Squares. NeuroImage, 3(3), 143–157. 10.1006/nimg.1996.0016

Mecca, A. P., McDonald, J. W., Michalak, H. R., Godek, T. A., Harris, J. E., Pugh, E. A., Kemp, E. C., Chen, M.-K., Salardini, A., Nabulsi, N. B., Lim, K., Huang, Y., Carson, R. E., Strittmatter, S. M., & van Dyck, C. H. (2020). PET imaging of mGluR5 in Alzheimer’s disease. Alzheimer’s Research & Therapy, 12(1), 15. 10.1186/s13195-020-0582-0

Mecca, A. P., Rogers, K., Jacobs, Z., McDonald, J. W., Michalak, H. R., DellaGioia, N., Zhao, W., Hillmer, A. T., Nabulsi, N., Lim, K., Ropchan, J., Huang, Y., Matuskey, D., Esterlis, I., Carson, R. E., & van Dyck, C. H. (2021). Effect of age on brain metabotropic glutamate receptor subtype 5 measured with [18F]FPEB PET. NeuroImage, 238, 118217. 10.1016/j.neuroimage.2021.118217

Mehta, K., Salo, T., Madison, T. J., Adebimpe, A., Bassett, D. S., Bertolero, M., Cieslak, M., Covitz, S., Houghton, A., Keller, A. S., Lundquist, J. T., Luo, A., Miranda-Dominguez, O., Nelson, S. M., Shafiei, G., Shanmugan, S., Shinohara, R. T., Smyser, C. D., Sydnor, V. J., … Satterthwaite, T. D. (2024). XCP-D: A robust pipeline for the post-processing of fMRI data. Imaging Neuroscience, 2, 1–26. 10.1162/imag_a_00257

Menon, V., & Uddin, L. Q. (2010). Saliency, switching, attention and control: A network model of insula function. Brain Structure and Function, 214(5), 655–667. 10.1007/s00429-010-0262-0

Meunier, D., Stamatakis, E. A., & Tyler, L. K. (2014). Age-related functional reorganization, structural changes, and preserved cognition. Neurobiology of Aging, 35(1), 42–54. 10.1016/j.neurobiolaging.2013.07.003

Morcom, A. M., & Johnson, W. (2015). Neural Reorganization and Compensation in Aging. Journal of Cognitive Neuroscience, 27(7), 1275–1285. 10.1162/jocn_a_00783

Neyman, S., & Manahan-Vaughan, D. (2008). Metabotropic glutamate receptor 1 (mGluR1) and 5 (mGluR5) regulate late phases of LTP and LTD in the hippocampal CA1 region in vitro. European Journal of Neuroscience, 27(6), 1345–1352. 10.1111/j.1460-9568.2008.06109.x

Nikolenko, V. N., Oganesyan, M. V., Rizaeva, N. A., Kudryashova, V. A., Nikitina, A. T., Pavliv, M. P., Shchedrina, M. A., Giller, D. B., Bulygin, K. V., & Sinelnikov, M. Y. (2020). Amygdala: Neuroanatomical and Morphophysiological Features in Terms of Neurological and Neurodegenerative Diseases. Brain Sciences, 10(8), 502. 10.3390/brainsci10080502

Pessoa, L. (2010). Emotion and cognition and the amygdala: From “what is it?” to “what’s to be done?” Neuropsychologia, 48(12), 3416–3429. 10.1016/j.neuropsychologia.2010.06.038

Pessoa, L. (2017). A Network Model of the Emotional Brain. Trends in Cognitive Sciences, 21(5), 357–371. 10.1016/j.tics.2017.03.002

Phelps, E. A., & LeDoux, J. E. (2005). Contributions of the Amygdala to Emotion Processing: From Animal Models to Human Behavior. Neuron, 48(2), 175–187. 10.1016/j.neuron.2005.09.025

Poldrack, R. A., Kittur, A., Kalar, D., Miller, E., Seppa, C., Gil, Y., Parker, D. S., Sabb, F. W., & Bilder, R. M. (2011). The Cognitive Atlas: Toward a Knowledge Foundation for Cognitive Neuroscience. Frontiers in Neuroinformatics, 5. 10.3389/fninf.2011.00017

Price, J. L. (2003). Comparative Aspects of Amygdala Connectivity. Annals of the New York Academy of Sciences, 985(1), 50–58. 10.1111/j.1749-6632.2003.tb07070.x

Qin, L., & Gao, J.-H. (2021). New avenues for functional neuroimaging: Ultra-high field MRI and OPM-MEG. Psychoradiology, 1(4), 165–171. 10.1093/psyrad/kkab014

Rabinak, C. A., Angstadt, M., Welsh, R. C., Kennedy, A., Lyubkin, M., Martis, B., & Phan, K. L. (2011). Altered Amygdala Resting-State Functional Connectivity in Post-Traumatic Stress Disorder. Frontiers in Psychiatry, 2. 10.3389/fpsyt.2011.00062

Radhakrishnan, R., Nabulsi, N., Gaiser, E., Gallezot, J.-D., Henry, S., Planeta, B., Lin, S., Ropchan, J., Williams, W., Morris, E., D’Souza, D. C., Huang, Y., Carson, R. E., & Matuskey, D. (2018). Age-Related Change in 5-HT6 Receptor Availability in Healthy Male Volunteers Measured with 11C-GSK215083 PET. Journal of Nuclear Medicine, 59(9), 1445–1450. 10.2967/jnumed.117.206516

Raven, J. C. (1941). Standardization of progressive matrices, 1938. British Journal of Medical Psychology, 19, 137–150. 10.1111/j.2044-8341.1941.tb00316.x

Raz, N., Rodrigue, K. M., Kennedy, K. M., Dahle, C., Head, D., & Acker, J. D. (2003). Differential age-related changes in the regional metencephalic volumes in humans: A 5-year follow-up. Neuroscience Letters, 349(3), 163–166. 10.1016/S0304-3940(03)00820-6

Reitan, R. M., & Wolfson, D. (2004). The Trail Making Test as an initial screening procedure for neuropsychological impairment in older children. Archives of Clinical Neuropsychology, 19(2), 281–288. 10.1016/S0887-6177(03)00042-8

Ritchey, M., & Cooper, R. A. (2020). Deconstructing the Posterior Medial Episodic Network. Trends in Cognitive Sciences, 24(6), 451–465. 10.1016/j.tics.2020.03.006

Ritchey, M., Montchal, M. E., Yonelinas, A. P., & Ranganath, C. (2015). Delay-dependent contributions of medial temporal lobe regions to episodic memory retrieval. eLife, 4, e05025. 10.7554/eLife.05025

Ronde, M., van der Zee, E. A., & Kas, M. J. H. (2024). Default mode network dynamics: An integrated neurocircuitry perspective on social dysfunction in human brain disorders. Neuroscience & Biobehavioral Reviews, 164, 105839. 10.1016/j.neubiorev.2024.105839

Sadaghiani, S., & D’Esposito, M. (2015). Functional Characterization of the Cingulo-Opercular Network in the Maintenance of Tonic Alertness. Cerebral Cortex, 25(9), 2763–2773. 10.1093/cercor/bhu072

Sakaki, M., Nga, L., & Mather, M. (2013). Amygdala Functional Connectivity with Medial Prefrontal Cortex at Rest Predicts the Positivity Effect in Older Adults’ Memory. Journal of Cognitive Neuroscience, 25(8), 1206–1224. 10.1162/jocn_a_00392

Sambuco, N. (2024). Cognition, emotion, and the default mode network. Brain and Cognition, 182, 106229. 10.1016/j.bandc.2024.106229

Sampson, P. D., Streissguth, A. P., Barr, H. M., & Bookstein, F. L. (1989). Neurobehavioral effects of prenatal alcohol: Part II. Partial Least Squares analysis. Neurotoxicology and Teratology, 11(5), 477–491. 10.1016/0892-0362(89)90025-1

Sanda, P., Hlinka, J., Berg, M. van den, Skoch, A., Bazhenov, M., Keliris, G. A., & Krishnan, G. P. (2024). Cholinergic modulation supports dynamic switching of resting state networks through selective DMN suppression. PLOS Computational Biology, 20(6), e1012099. 10.1371/journal.pcbi.1012099

Schaefer, A., Kong, R., Gordon, E. M., Laumann, T. O., Zuo, X.-N., Holmes, A. J., Eickhoff, S. B., & Yeo, B. T. T. (2018). Local-Global Parcellation of the Human Cerebral Cortex from Intrinsic Functional Connectivity MRI. Cerebral Cortex, 28(9), 3095–3114. 10.1093/cercor/bhx179

Schroeder, R. W., Twumasi-Ankrah, P., Baade, L. E., & Marshall, P. S. (2012). Reliable Digit Span: A Systematic Review and Cross-Validation Study. Assessment, 19(1), 21–30. 10.1177/1073191111428764

Schweighofer, N., Bertin, M., Shishida, K., Okamoto, Y., Tanaka, S. C., Yamawaki, S., & Doya, K. (2008). Low-Serotonin Levels Increase Delayed Reward Discounting in Humans. Journal of Neuroscience, 28(17), 4528–4532. 10.1523/JNEUROSCI.4982-07.2008

Seeley, W. W. (2019a). The Salience Network: A Neural System for Perceiving and Responding to Homeostatic Demands. Journal of Neuroscience, 39(50), 9878–9882. 10.1523/JNEUROSCI.1138-17.2019

Seeley, W. W. (2019b). The Salience Network: A Neural System for Perceiving and Responding to Homeostatic Demands. Journal of Neuroscience, 39(50), 9878–9882. 10.1523/JNEUROSCI.1138-17.2019

Sengupta, A., Bocchio, M., Bannerman, D. M., Sharp, T., & Capogna, M. (2017). Control of Amygdala Circuits by 5-HT Neurons via 5-HT and Glutamate Cotransmission. Journal of Neuroscience, 37(7), 1785–1796. 10.1523/JNEUROSCI.2238-16.2016

Singh, S., Malo, P. K., Stezin, A., Mensegere, A. L., & Issac, T. G. (2024). Alteration in amygdala subfield volumes and their association with cognition in mild cognitive impairment. Journal of Neurology, 271(8), 5460–5467. 10.1007/s00415-024-12500-3

Siwek, A., Marcinkowska, M., Głuch-Lutwin, M., Mordyl, B., Wolak, M., Jastrzębska-Więsek, M., Wilczyńska-Zawal, N., Wyska, E., Szafrańska, K., Karcz, T., Ostrowska, O., Bucki, A., & Kołaczkowski, M. (2024). Dual 5-HT6/SERT ligands for mitigating neuropsychiatric symptoms of dementia exerting neuroprotection against amyloid-β toxicity, memory preservation, and antidepressant-like properties. European Journal of Medicinal Chemistry, 275, 116601. 10.1016/j.ejmech.2024.116601

Sladky, R., Baldinger, P., Kranz, G. S., Tröstl, J., Höflich, A., Lanzenberger, R., Moser, E., & Windischberger, C. (2013). High-resolution functional MRI of the human amygdala at 7 T. European Journal of Radiology, 82(5), 728–733. 10.1016/j.ejrad.2011.09.025

Song, J., Birn, R. M., Boly, M., Meier, T. B., Nair, V. A., Meyerand, M. E., & Prabhakaran, V. (2014). Age-Related Reorganizational Changes in Modularity and Functional Connectivity of Human Brain Networks. Brain Connectivity, 4(9), 662–676. 10.1089/brain.2014.0286

Spaniol, J., & Grady, C. (2012). Aging and the neural correlates of source memory: Over-recruitment and functional reorganization. Neurobiology of Aging, 33(2), 425.e3-425.e18. 10.1016/j.neurobiolaging.2010.10.005

Spreng, R. N., Stevens, W. D., Chamberlain, J. P., Gilmore, A. W., & Schacter, D. L. (2010). Default network activity, coupled with the frontoparietal control network, supports goal-directed cognition. NeuroImage, 53(1), 303–317. 10.1016/j.neuroimage.2010.06.016

Spreng, R. N., & Turner, G. R. (2019). The Shifting Architecture of Cognition and Brain Function in Older Adulthood. Perspectives on Psychological Science, 14(4), 523–542. 10.1177/1745691619827511

St. Jacques, P., Dolcos, F., & Cabeza, R. (2010). Effects of aging on functional connectivity of the amygdala during negative evaluation: A network analysis of fMRI data. Neurobiology of Aging, 31(2), 315–327. 10.1016/j.neurobiolaging.2008.03.012

Straube, T., Preissler, S., Lipka, J., Hewig, J., Mentzel, H.-J., & Miltner, W. H. R. (2010). Neural representation of anxiety and personality during exposure to anxiety-provoking and neutral scenes from scary movies. Human Brain Mapping, 31(1), 36–47. 10.1002/hbm.20843

Thomas Yeo, B. T., Krienen, F. M., Sepulcre, J., Sabuncu, M. R., Lashkari, D., Hollinshead, M., Roffman, J. L., Smoller, J. W., Zöllei, L., Polimeni, J. R., Fischl, B., Liu, H., & Buckner, R. L. (2011). The organization of the human cerebral cortex estimated by intrinsic functional connectivity. Journal of Neurophysiology, 106(3), 1125–1165. 10.1152/jn.00338.2011

Tian, Y., Margulies, D. S., Breakspear, M., & Zalesky, A. (2020). Topographic organization of the human subcortex unveiled with functional connectivity gradients. Nat. Neurosci, 23, 1421–1432.

Ueno, D., Matsuoka, T., Kato, Y., Ayani, N., Maeda, S., Takeda, M., & Narumoto, J. (2020). Individual Differences in Interoceptive Accuracy Are Correlated With Salience Network Connectivity in Older Adults. Frontiers in Aging Neuroscience, 12. 10.3389/fnagi.2020.592002

Vossel, S., Geng, J. J., & Fink, G. R. (2014). Dorsal and Ventral Attention Systems: Distinct Neural Circuits but Collaborative Roles. The Neuroscientist, 20(2), 150–159. 10.1177/1073858413494269

Walhovd, K. B., Westlye, L. T., Amlien, I., Espeseth, T., Reinvang, I., Raz, N., Agartz, I., Salat, D. H., Greve, D. N., Fischl, B., Dale, A. M., & Fjell, A. M. (2011). Consistent neuroanatomical age-related volume differences across multiple samples. Neurobiology of Aging, 32(5), 916–932. 10.1016/j.neurobiolaging.2009.05.013

Wang, J., Sun, L., Chen, L., Sun, J., Xie, Y., Tian, D., Gao, L., Zhang, D., Xia, M., & Wu, T. (2023). Common and distinct roles of amygdala subregional functional connectivity in non-motor symptoms of Parkinson’s disease. Npj Parkinson’s Disease, 9(1), 28. 10.1038/s41531-023-00469-1

Werneke, U., Goldberg, D. P., Yalcin, I., & Üstün, B. T. (2000). The stability of the factor structure of the General Health Questionnaire. Psychological Medicine, 30(4), 823–829. 10.1017/S0033291799002287

Wesołowska, A. (2010). Potential role of the 5-HT6 receptor in depression and anxiety: An overview of preclinical data. Pharmacological Reports, 62(4), 564–577. 10.1016/S1734-1140(10)70315-7

Wiebels, K., Waldie, K. E., Roberts, R. P., & Park, H. R. P. (2016). Identifying grey matter changes in schizotypy using partial least squares correlation. Cortex, 81, 137–150. 10.1016/j.cortex.2016.04.011

Worbe, Y., Savulich, G., Voon, V., Fernandez-Egea, E., & Robbins, T. W. (2014). Serotonin Depletion Induces ‘Waiting Impulsivity’ on the Human Four-Choice Serial Reaction Time Task: Cross-Species Translational Significance. Neuropsychopharmacology, 39(6), 1519–1526. 10.1038/npp.2013.351

Yarkoni, T., Poldrack, R. A., Nichols, T. E., Van Essen, D. C., & Wager, T. D. (2011). Large-scale automated synthesis of human functional neuroimaging data. Nature Methods, 8(8), 665–670. 10.1038/nmeth.1635

Ye, S., Bätz, L. R., Jain, A., Salami, A., & Ziaei, M. (2024). Resilience-driven neural synchrony during naturalistic movie watching (p. 2023.10.12.562025). bioRxiv. 10.1101/2023.10.12.562025

Ye, S., Dave, A., Salami, A., & Ziaei, M. (2025). Frontoparietal functional dedifferentiation during naturalistic movie watching among older adults at risk of emotional vulnerability. Neurobiology of Aging, 156, 150–162. 10.1016/j.neurobiolaging.2025.09.004

Zhang, Y., Zhou, W., Huang, J., Hong, B., & Wang, X. (2022). Neural correlates of perceived emotions in human insula and amygdala for auditory emotion recognition. NeuroImage, 260, 119502. 10.1016/j.neuroimage.2022.119502

Zhou, F., & Becker, B. (2025). The default mode network and emotion—Dual roles of the medial prefrontal cortex in emotional experience and regulation. Current Opinion in Behavioral Sciences, 66, 101613. 10.1016/j.cobeha.2025.101613

Zhou, Y., & Danbolt, N. C. (2014). Glutamate as a neurotransmitter in the healthy brain. Journal of Neural Transmission, 121(8), 799–817. 10.1007/s00702-014-1180-8

Ziaei, M., Bonyadi, M. R., & Reutens, D. C. (2021). Age-related differences in structural and functional prefrontal networks during a logical reasoning task. Brain Imaging and Behavior, 15(2), 1085–1102. 10.1007/s11682-020-00315-5

Ziaei, M., Burianová, H., von Hippel, W., Ebner, N. C., Phillips, L. H., & Henry, J. D. (2016). The impact of aging on the neural networks involved in gaze and emotional processing. Neurobiology of Aging, 48, 182–194. 10.1016/j.neurobiolaging.2016.08.026

Ziaei, M., Oestreich, L., Persson, J., Reutens, D. C., & Ebner, N. C. (2022). Neural correlates of affective empathy in aging: A multimodal imaging and multivariate approach: Abbreviated title: Multimodal and multivariate approach to empathy. Aging, Neuropsychology, and Cognition, 29(3), 577–598. 10.1080/13825585.2022.2036684

Ziaei, M., Salami, A., & Persson, J. (2017). Age-related alterations in functional connectivity patterns during working memory encoding of emotional items. Neuropsychologia, 94, 1–12. 10.1016/j.neuropsychologia.2016.11.012

Zöller, D., Schaer, M., Scariati, E., Padula, M. C., Eliez, S., & Van De Ville, D. (2017). Disentangling resting-state BOLD variability and PCC functional connectivity in 22q11.2 deletion syndrome. NeuroImage, 149, 85–97. 10.1016/j.neuroimage.2017.01.064

