## supplementary for "Functional connectivity profile of the amygdala subfields associates with emotional well-being in aging"

**This file includes:**

Supplementary method

Tables S1

Figures S1 to S5

**Imaging data preprocessing**

Results included in this manuscript come from preprocessing performed using *fMRIPrep* 22.0.2 (Esteban, Markiewicz, et al. (2018); Esteban, Blair, et al. (2018); RRID:SCR_016216), which is based on *Nipype* 1.8.5 (K. Gorgolewski et al. (2011); K. J. Gorgolewski et al. (2018); RRID:SCR_002502).

**Preprocessing of B0 inhomogeneity mappings**

A total of 2 fieldmaps were found available within the input BIDS structure. A B0-nonuniformity map (or fieldmap) was estimated based on two (or more) echo-planar imaging (EPI) references with topup (Andersson, Skare, and Ashburner (2003); FSL 6.0.5.1:57b01774).

**Anatomical data preprocessing**

A total of 1 T1-weighted (T1w) images were found within the input BIDS dataset. The T1-weighted (T1w) image was corrected for intensity non-uniformity (INU) with N4BiasFieldCorrection (Tustison et al. 2010), distributed with ANTs 2.3.3 (Avants et al. 2008, RRID:SCR_004757), and used as T1w-reference throughout the workflow. The T1w-reference was then skull-stripped with a *Nipype* implementation of the antsBrainExtraction.sh workflow (from ANTs), using OASIS30ANTs as target template. Brain tissue segmentation of cerebrospinal fluid (CSF), white-matter (WM) and gray-matter (GM) was performed on the brain-extracted T1w using fast (FSL 6.0.5.1:57b01774, RRID:SCR_002823, Zhang, Brady, and Smith 2001). Brain surfaces were reconstructed using recon-all (FreeSurfer 7.2.0, RRID:SCR_001847, Dale, Fischl, and Sereno 1999), and the brain mask estimated previously was refined with a custom variation of the method to reconcile ANTs-derived and FreeSurfer-derived segmentations of the cortical gray-matter of Mindboggle (RRID:SCR_002438, Klein et al. 2017). Volume-based spatial normalization to one standard space (MNI152NLin2009cAsym) was performed through nonlinear registration with antsRegistration (ANTs 2.3.3), using brain-extracted versions of both T1w reference and the T1w template. The following template was selected for spatial normalization: *ICBM 152 Nonlinear Asymmetrical template version 2009c* [Fonov et al. (2009), RRID:SCR_008796; TemplateFlow ID: MNI152NLin2009cAsym].

**Functional data preprocessing**

For each of the 2 BOLD runs found per subject (across all tasks and sessions), the following preprocessing was performed. First, a reference volume and its skull-stripped version were generated using a custom methodology of *fMRIPrep*. Head-motion parameters with respect to the BOLD reference (transformation matrices, and six corresponding rotation and translation parameters) are estimated before any spatiotemporal filtering using mcflirt (FSL 6.0.5.1:57b01774, Jenkinson et al. 2002). BOLD runs were slice-time corrected to 0.978s (0.5 of slice acquisition range 0s-1.96s) using 3dTshift from AFNI (Cox and Hyde 1997, RRID:SCR_005927). The BOLD time-series (including slice-timing correction when applied) were resampled onto their original, native space by applying the transforms to correct for head-motion. These resampled BOLD time-series will be referred to as *preprocessed BOLD in original space*, or just *preprocessed BOLD*. The BOLD reference was then co-registered to the T1w reference using bbregister (FreeSurfer) which implements boundary-based registration (Greve and Fischl 2009). Co-registration was configured with six degrees of freedom. Several confounding time-series were calculated based on the *preprocessed BOLD*: framewise displacement (FD), DVARS and three region-wise global signals. FD was computed using two formulations following Power (absolute sum of relative motions, Power et al. (2014)) and Jenkinson (relative root mean square displacement between affines, Jenkinson et al. (2002)). FD and DVARS are calculated for each functional run, both using their implementations in *Nipype* (following the definitions by Power et al. 2014). The three global signals are extracted within the CSF, the WM, and the whole-brain masks. Additionally, a set of physiological regressors were extracted to allow for component-based noise correction (*CompCor*, Behzadi et al. 2007). Principal components are estimated after high-pass filtering the *preprocessed BOLD* time-series (using a discrete cosine filter with 128s cut-off) for the two *CompCor* variants: temporal (tCompCor) and anatomical (aCompCor). tCompCor components are then calculated from the top 2% variable voxels within the brain mask. For aCompCor, three probabilistic masks (CSF, WM and combined CSF+WM) are generated in anatomical space. The implementation differs from that of Behzadi et al. in that instead of eroding the masks by 2 pixels on BOLD space, a mask of pixels that likely contain a volume fraction of GM is subtracted from the aCompCor masks. This mask is obtained by dilating a GM mask extracted from the FreeSurfer’s *aseg* segmentation, and it ensures components are not extracted from voxels containing a minimal fraction of GM. Finally, these masks are resampled into BOLD space and binarized by thresholding at 0.99 (as in the original implementation). Components are also calculated separately within the WM and CSF masks. For each CompCor decomposition, the *k* components with the largest singular values are retained, such that the retained components’ time series are sufficient to explain 50 percent of variance across the nuisance mask (CSF, WM, combined, or temporal). The remaining components are dropped from consideration. The head-motion estimates calculated in the correction step were also placed within the corresponding confounds file. The confound time series derived from head motion estimates and global signals were expanded with the inclusion of temporal derivatives and quadratic terms for each (Satterthwaite et al. 2013). Frames that exceeded a threshold of 0.5 mm FD or 1.5 standardized DVARS were annotated as motion outliers. Additional nuisance timeseries are calculated by means of principal components analysis of the signal found within a thin band (*crown*) of voxels around the edge of the brain, as proposed by (Patriat, Reynolds, and Birn 2017). The BOLD time-series were resampled into standard space, generating a *preprocessed BOLD run in MNI152NLin2009cAsym space*. First, a reference volume and its skull-stripped version were generated using a custom methodology of *fMRIPrep*. All resamplings can be performed with *a single interpolation step* by composing all the pertinent transformations (i.e. head-motion transform matrices, susceptibility distortion correction when available, and co-registrations to anatomical and output spaces). Gridded (volumetric) resamplings were performed using antsApplyTransforms (ANTs), configured with Lanczos interpolation to minimize the smoothing effects of other kernels (Lanczos 1964). Non-gridded (surface) resamplings were performed using mri_vol2surf (FreeSurfer).

Many internal operations of *fMRIPrep* use *Nilearn* 0.9.1 (Abraham et al. 2014, RRID:SCR_001362), mostly within the functional processing workflow. For more details of the pipeline, see [the section corresponding to workflows in *fMRIPrep*’s documentation](https://fmriprep.readthedocs.io/en/latest/workflows.html).

Copyright Waiver

The above boilerplate text was automatically generated by fMRIPrep with the express intention that users should copy and paste this text into their manuscripts *unchanged*. It is released under the [CC0](https://creativecommons.org/publicdomain/zero/1.0/) license.

References

Abraham, Alexandre, Fabian Pedregosa, Michael Eickenberg, Philippe Gervais, Andreas Mueller, Jean Kossaifi, Alexandre Gramfort, Bertrand Thirion, and Gael Varoquaux. 2014. “Machine Learning for Neuroimaging with Scikit-Learn.” *Frontiers in Neuroinformatics* 8. <https://doi.org/10.3389/fninf.2014.00014>.

Avants, B. B., C. L. Epstein, M. Grossman, and J. C. Gee. 2008. “Symmetric Diffeomorphic Image Registration with Cross-Correlation: Evaluating Automated Labeling of Elderly and Neurodegenerative Brain.” *Medical Image Analysis* 12 (1): 26–41. <https://doi.org/10.1016/j.media.2007.06.004>.

Behzadi, Yashar, Khaled Restom, Joy Liau, and Thomas T. Liu. 2007. “A Component Based Noise Correction Method (CompCor) for BOLD and Perfusion Based fMRI.” *NeuroImage* 37 (1): 90–101. <https://doi.org/10.1016/j.neuroimage.2007.04.042>.

Cox, Robert W., and James S. Hyde. 1997. “Software Tools for Analysis and Visualization of fMRI Data.” *NMR in Biomedicine* 10 (4-5): 171–78. [https://doi.org/10.1002/(SICI)1099-1492(199706/08)10:4/5<171::AID-NBM453>3.0.CO;2-L](https://doi.org/10.1002/(SICI)1099-1492(199706/08)10:4/5%3c171::AID-NBM453%3e3.0.CO;2-L).

Dale, Anders M., Bruce Fischl, and Martin I. Sereno. 1999. “Cortical Surface-Based Analysis: I. Segmentation and Surface Reconstruction.” *NeuroImage* 9 (2): 179–94. <https://doi.org/10.1006/nimg.1998.0395>.

Esteban, Oscar, Ross Blair, Christopher J. Markiewicz, Shoshana L. Berleant, Craig Moodie, Feilong Ma, Ayse Ilkay Isik, et al. 2018. “fMRIPrep 22.0.2.” *Software*. <https://doi.org/10.5281/zenodo.852659>.

Esteban, Oscar, Christopher Markiewicz, Ross W Blair, Craig Moodie, Ayse Ilkay Isik, Asier Erramuzpe Aliaga, James Kent, et al. 2018. “fMRIPrep: A Robust Preprocessing Pipeline for Functional MRI.” *Nature Methods*. <https://doi.org/10.1038/s41592-018-0235-4>.

Fonov, VS, AC Evans, RC McKinstry, CR Almli, and DL Collins. 2009. “Unbiased Nonlinear Average Age-Appropriate Brain Templates from Birth to Adulthood.” *NeuroImage* 47, Supplement 1: S102. <https://doi.org/10.1016/S1053-8119(09)70884-5>.

Gorgolewski, K., C. D. Burns, C. Madison, D. Clark, Y. O. Halchenko, M. L. Waskom, and S. Ghosh. 2011. “Nipype: A Flexible, Lightweight and Extensible Neuroimaging Data Processing Framework in Python.” *Frontiers in Neuroinformatics* 5: 13. <https://doi.org/10.3389/fninf.2011.00013>.

Gorgolewski, Krzysztof J., Oscar Esteban, Christopher J. Markiewicz, Erik Ziegler, David Gage Ellis, Michael Philipp Notter, Dorota Jarecka, et al. 2018. “Nipype.” *Software*. <https://doi.org/10.5281/zenodo.596855>.

Greve, Douglas N, and Bruce Fischl. 2009. “Accurate and Robust Brain Image Alignment Using Boundary-Based Registration.” *NeuroImage* 48 (1): 63–72. <https://doi.org/10.1016/j.neuroimage.2009.06.060>.

Jenkinson, Mark, Peter Bannister, Michael Brady, and Stephen Smith. 2002. “Improved Optimization for the Robust and Accurate Linear Registration and Motion Correction of Brain Images.” *NeuroImage* 17 (2): 825–41. <https://doi.org/10.1006/nimg.2002.1132>.

Klein, Arno, Satrajit S. Ghosh, Forrest S. Bao, Joachim Giard, Yrjö Häme, Eliezer Stavsky, Noah Lee, et al. 2017. “Mindboggling Morphometry of Human Brains.” *PLOS Computational Biology* 13 (2): e1005350. <https://doi.org/10.1371/journal.pcbi.1005350>.

Lanczos, C. 1964. “Evaluation of Noisy Data.” *Journal of the Society for Industrial and Applied Mathematics Series B Numerical Analysis* 1 (1): 76–85. <https://doi.org/10.1137/0701007>.

Patriat, Rémi, Richard C. Reynolds, and Rasmus M. Birn. 2017. “An Improved Model of Motion-Related Signal Changes in fMRI.” *NeuroImage* 144, Part A (January): 74–82. <https://doi.org/10.1016/j.neuroimage.2016.08.051>.

Power, Jonathan D., Anish Mitra, Timothy O. Laumann, Abraham Z. Snyder, Bradley L. Schlaggar, and Steven E. Petersen. 2014. “Methods to Detect, Characterize, and Remove Motion Artifact in Resting State fMRI.” *NeuroImage* 84 (Supplement C): 320–41. <https://doi.org/10.1016/j.neuroimage.2013.08.048>.

Satterthwaite, Theodore D., Mark A. Elliott, Raphael T. Gerraty, Kosha Ruparel, James Loughead, Monica E. Calkins, Simon B. Eickhoff, et al. 2013. “An improved framework for confound regression and filtering for control of motion artifact in the preprocessing of resting-state functional connectivity data.” *NeuroImage* 64 (1): 240–56. <https://doi.org/10.1016/j.neuroimage.2012.08.052>.

Tustison, N. J., B. B. Avants, P. A. Cook, Y. Zheng, A. Egan, P. A. Yushkevich, and J. C. Gee. 2010. “N4itk: Improved N3 Bias Correction.” *IEEE Transactions on Medical Imaging* 29 (6): 1310–20. <https://doi.org/10.1109/TMI.2010.2046908>.

Zhang, Y., M. Brady, and S. Smith. 2001. “Segmentation of Brain MR Images Through a Hidden Markov Random Field Model and the Expectation-Maximization Algorithm.” *IEEE Transactions on Medical Imaging* 20 (1): 45–57. <https://doi.org/10.1109/42.906424>.

**Post-processing of fmriprep outputs**

The eXtensible Connectivity Pipeline (XCP) (Ciric et al. 2018; Satterthwaite et al. 2013) was used to post-process the outputs of fmriprep version 22.0.2 (Esteban et al. 2019, 2020, RRID:SCR_016216). XCP was built with *Nipype* 1.8.5 (Gorgolewski et al. 2011, RRID:SCR_002502). For each of the two BOLD series found per subject (across all tasks and sessions), the following post-processing was performed. First, the first three of both the BOLD data and nuisance regressors were discarded, then outlier detection was performed. In order to identify high-motion outlier volumes, framewise displacement was calculated using the formula from Power et al. (2014), with a head radius of 50 mm. Volumes with framewise displacement greater than 0.2 mm were flagged as outliers and excluded from nuisance regression (Power et al. 2014).

Before nuisance regression, but after censoring, the BOLD data were mean-centered and linearly detrended. The top 5 principal aCompCor components from WM and CSF compartments were selected as nuisance regressors. Additionally, the six motion parameters and their temporal derivatives were added as confounds. (Ciric et al. 2017; Satterthwaite et al. 2013). These nuisance regressors were regressed from the BOLD data using linear regression - as implemented in Scikit-Learn 1.1.3 (Pedregosa et al. 2011). Any volumes censored earlier in the workflow were then interpolated in the residual time series produced by the regression. The interpolated timeseries were then band-pass filtered to retain signals within the 0.009-0.08 Hz frequency band. The processed BOLD was smoothed using Nilearn with a gaussian kernel size of 4.0 mm (FWHM).

Many internal operations of *XCP* use *TemplateFlow* version 0.8.1 (Ciric et al. 2022), *Nibabel* version 4.0.2 (Brett et al. 2022), *numpy* version 1.18.5 (Harris et al. 2020), and *scipy* version 1.9.3 (Virtanen et al. 2020). For more details, see the *xcp_d* website https://xcp-d.readthedocs.io.

Copyright Waiver

The above methods description text was automatically generated by *XCP* with the express intention that users should copy and paste this text into their manuscripts *unchanged*. It is released under the [CC0](https://creativecommons.org/publicdomain/zero/1.0/) license.

References

Abraham, Alexandre, Fabian Pedregosa, Michael Eickenberg, Philippe Gervais, Andreas Mueller, Jean Kossaifi, Alexandre Gramfort, Bertrand Thirion, and Gael Varoquaux. 2014. “Machine Learning for Neuroimaging with Scikit-Learn.” *Frontiers in Neuroinformatics*. <https://doi.org/10.3389/fninf.2014.00014>.

Brett, Matthew, Christopher J. Markiewicz, Michael Hanke, Marc-Alexandre Côté, Ben Cipollini, Paul McCarthy, Dorota Jarecka, et al. 2022. *Nipy/Nibabel:* (version 4.0.0). Zenodo. <https://doi.org/10.5281/zenodo.591597>.

Ciric, Rastko, Adon F. G. Rosen, Guray Erus, Matthew Cieslak, Azeez Adebimpe, Philip A. Cook, Danielle S. Bassett, Christos Davatzikos, Daniel H. Wolf, and Theodore D. Satterthwaite. 2018. “Mitigating Head Motion Artifact in Functional Connectivity MRI.” *Nature Protocols* 13 (12): 2801–26. <https://doi.org/10.1038/s41596-018-0065-y>.

Ciric, Rastko, William H Thompson, Romy Lorenz, Mathias Goncalves, Eilidh MacNicol, Christopher J Markiewicz, Yaroslav O Halchenko, et al. 2022. “TemplateFlow: FAIR-Sharing of Multi-Scale, Multi-Species Brain Models.” *bioRxiv*. Cold Spring Harbor Laboratory, 2021–02. <https://doi.org/10.1101/2021.02.10.430678>.

Ciric, Rastko, Daniel H. Wolf, Jonathan D. Power, David R. Roalf, Graham Baum, Kosha Ruparel, Russell T. Shinohara, et al. 2017. “Benchmarking of Participant-Level Confound Regression Strategies for the Control of Motion Artifact in Studies of Functional Connectivity.” *NeuroImage* 154 (July): 174–87. <https://doi.org/10.1016/j.neuroimage.2017.03.020>.

Cox, Robert, and James Hyd. 1997. “Software Tools for Analysis and Visualization of fMRI Data.” *NMR in Biomedicine* 10 (4-5): 171–78. [https://doi.org/10.1002/(sici)1099-1492(199706/08)10:4/5<171::aid-nbm453>3.0.co;2-l](https://doi.org/10.1002/(sici)1099-1492(199706/08)10:4/5%3c171::aid-nbm453%3e3.0.co;2-l).

Esteban, Oscar, Rastko Ciric, Karolina Finc, Ross W Blair, Christopher J Markiewicz, Craig A Moodie, James D Kent, et al. 2020. “Analysis of Task-Based Functional Mri Data Preprocessed with fMRIPrep.” *Nature Protocols* 15 (7). Nature Publishing Group: 2186–2202. <https://doi.org/10.1038/s41596-020-0327-3>.

Esteban, Oscar, Christopher J Markiewicz, Ross W Blair, Craig A Moodie, A Ilkay Isik, Asier Erramuzpe, James D Kent, et al. 2019. “FMRIPrep: A Robust Preprocessing Pipeline for Functional Mri.” *Nature Methods* 16 (1). Nature Publishing Group: 111–16. <https://doi.org/10.1038/s41592-018-0235-4>.

Glasser, Matthew F., Timothy S. Coalson, Emma C. Robinson, Carl D. Hacker, John Harwell, Essa Yacoub, Kamil Ugurbil, et al. 2016. “A Multi-Modal Parcellation of Human Cerebral Cortex.” *Nature* 536 (7615): 171–78. <https://doi.org/10.1038/nature18933>.

Gordon, Evan M., Timothy O. Laumann, Babatunde Adeyemo, Jeremy F. Huckins, William M. Kelley, and Steven E. Petersen. 2016. “Generation and Evaluation of a Cortical Area Parcellation from Resting-State Correlations.” *Cerebral Cortex* 26 (1): 288–303. <https://doi.org/10.1093/cercor/bhu239>.

Gorgolewski, Krzysztof, Christopher D. Burns, Cindee Madison, Dav Clark, Yaroslav O. Halchenko, Michael L. Waskom, and Satrajit S. Ghosh. 2011. “Nipype: A Flexible, Lightweight and Extensible Neuroimaging Data Processing Framework in Python.” *Frontiers in Neuroinformatics* 5. <https://doi.org/10.3389/fninf.2011.00013>.

Harris, Charles R., Jarrod K. Millman, Stéfan J. van der Walt, Ralf Gommers, Pauli Virtanen, David Cournapeau, Eric Wieser, et al. 2020. “Array Programming with NumPy.” *Nature* 585 (7825): 357–62. <https://doi.org/10.1038/s41586-020-2649-2>.

Pedregosa, Fabian, Gael Varoquaux, Alexandre Gramfort, Vincent Michel, Bertrand Thirion, Olivier Grisel, Mathieu Blondel, et al. 2011. “Scikit-Learn: Machine Learning in Python.” *Journal of Machine Learning Research* 12: 2825–30.

Power, Jonathan D., Anish Mitra, Timothy O. Laumann, Abraham Z. Snyder, Bradley L. Schlaggar, and Steven E. Petersen. 2014. “Methods to Detect, Characterize, and Remove Motion Artifact in Resting State fMRI.” *NeuroImage* 84 (January): 320–41. <https://doi.org/10.1016/j.neuroimage.2013.08.048>.

Satterthwaite, Theodore D., Mark A. Elliott, Raphael T. Gerraty, Kosha Ruparel, James Loughead, Monica E. Calkins, Simon B. Eickhoff, et al. 2013. “An Improved Framework for Confound Regression and Filtering for Control of Motion Artifact in the Preprocessing of Resting-State Functional Connectivity Data.” *NeuroImage* 64 (January): 240–56. <https://doi.org/10.1016/j.neuroimage.2012.08.052>.

Schaefer, Alexander, Ru Kong, Evan M. Gordon, Timothy O. Laumann, Xi-Nian Zuo, Avram J. Holmes, Simon B. Eickhoff, and B. T. Thomas Yeo. 2018. “Local-Global Parcellation of the Human Cerebral Cortex from Intrinsic Functional Connectivity MRI.” *Cerebral Cortex (New York, N.Y.: 1991)* 28 (9): 3095–3114. <https://doi.org/10.1093/cercor/bhx179>.

Tian, Ye, Daniel S Margulies, Michael Breakspear, and Andrew Zalesky. 2020. “Topographic Organization of the Human Subcortex Unveiled with Functional Connectivity Gradients.” *Nature Neuroscience* 23 (11). Nature Publishing Group: 1421–32. <https://doi.org/10.1038/s41593-020-00711-6>.

Virtanen, Pauli, Ralf Gommers, Travis E. Oliphant, Matt Haberland, Tyler Reddy, David Cournapeau, Evgeni Burovski, et al. 2020. “SciPy 1.0: Fundamental Algorithms for Scientific Computing in Python.” *Nature Methods* 17 (3): 261–72. <https://doi.org/10.1038/s41592-019-0686-2>.

Zou, Qi-Hong, Chao-Zhe Zhu, Yihong Yang, Xi-Nian Zuo, Xiang-Yu Long, Qing-Jiu Cao, Yu-Feng Wang, and Yu-Feng Zang. 2008. “An Improved Approach to Detection of Amplitude of Low-Frequency Fluctuation (ALFF) for Resting-State fMRI: Fractional ALFF.” *Journal of Neuroscience Methods* 172 (1): 137–41. <https://doi.org/10.1016/j.jneumeth.2008.04.012>.

**Table S1. Variance explained and permutation *p*-values for latent variables (LVs) identified in the PLS analyses.**

|  | **Variance explained (permutation *p*-value)** | | | |
| --- | --- | --- | --- | --- |
|  | **LV1** | **LV2** | **LV3** | **LV4** |
| ***Neutral movie*** |  |  |  |  |
| BLA | 32.5% (0.136) | 27.0% (0.586) | 25.1% (0.404) | 15.5% (0.901) |
| CMA | 39.3% (0.128) | 26.0% (0.486) | 19.6% (0.849) | 15.1% (0.653) |
| SFA | 43.3% (0.064) | 22.8% (0.455) | 20.0% (0.862) | 13.9% (0.832) |
| Amygdala | 34.0% (0.287) | 25.7% (0.396) | 25.0% (0.429) | 15.4% (0.762) |
| ***Negative movie*** |  |  |  |  |
| BLA | 56.0% (0.005) | 18.3% (0.368) | 16.0% (0.793) | 9.7% (0.990) |
| CMA | 60.6% (0.001) | 17.7% (0.324) | 12.1% (0.988) | 9.6% (0.922) |
| SFA | 54.6% (0.006) | 18.4% (0.486) | 15.2% (0.934) | 11.7% (0.908) |
| Amygdala | 53.9% (0.004) | 19.7% (0.364) | 16.6% (0.711) | 9.8% (0.986) |

Note: BLA = basolateral amygdala; CMA = centromedial amygdala; SFA = superficial amygdala.

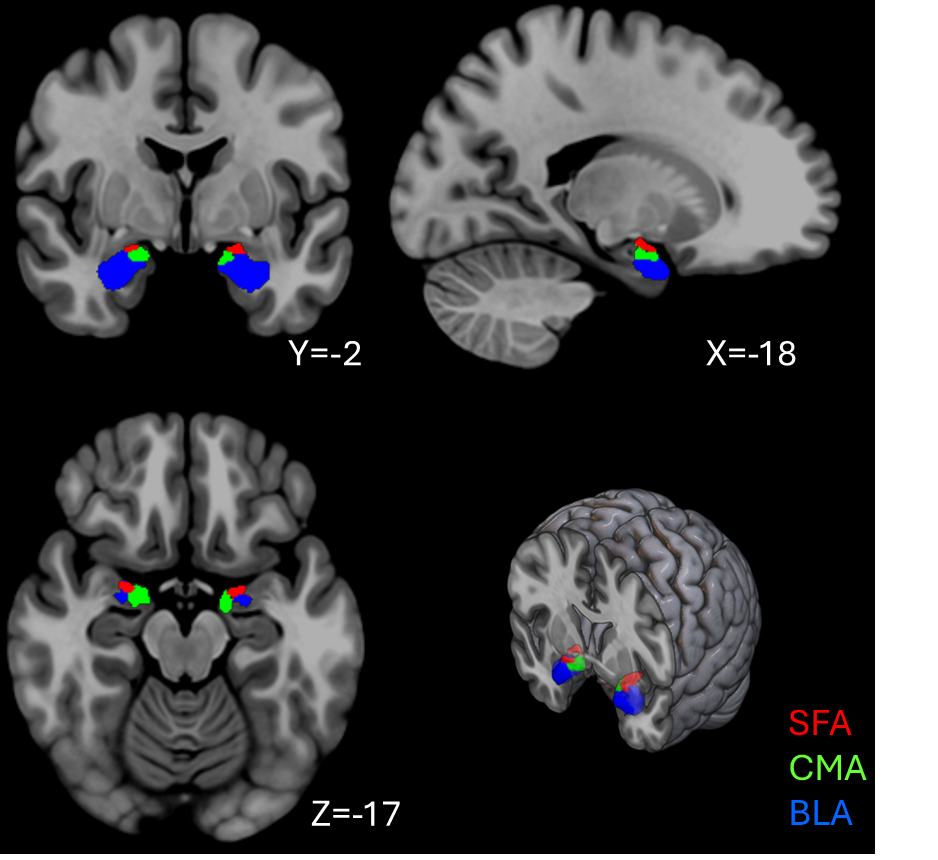

**Figure S1. Anatomical localization of amygdala subregions.** The three amygdala subregions used as seed regions for functional connectivity analyses are displayed on the MNI152 template brain. The superficial (SFA, red), centromedial (CMA, green), and basolateral (BLA, blue) subregions are shown in 2D and 3D views. Subregional boundaries were defined using the Tian subcortical atlas.

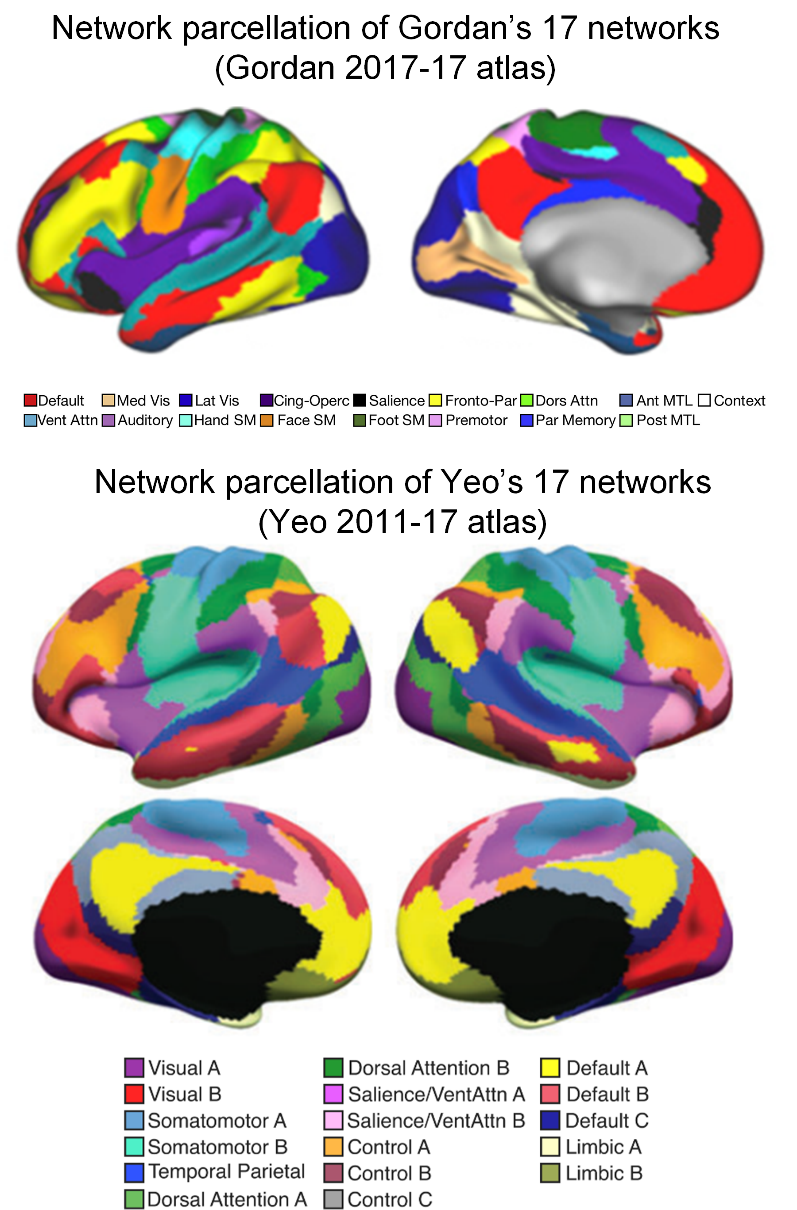

**Figure S2. Cortical network parcellations applied in the network correspondence analyses.** Surface maps illustrate the 17 canonical functional networks from the Gordon 2017–17 atlas (Top) and the Yeo 2011–17 atlas (Bottom). Default= default mode network; Med Vis=medial visual network; Lat Vis=lateral visual network. Cing-Operc=cingulo-opercular network; Fronto-Par=fronto-parietal network; Dors Attn=dorsal attention network; Ant MTL=anterior medial temporal network; Context=contextual association network; Vent Attn=ventral attention network; Hand SM=hand somatomotor network; Face SM=face somatomotor network; Foot SM=foot somatomotor network; Par Memory=parietal memory; Post MTL=posterior medial temporal network.

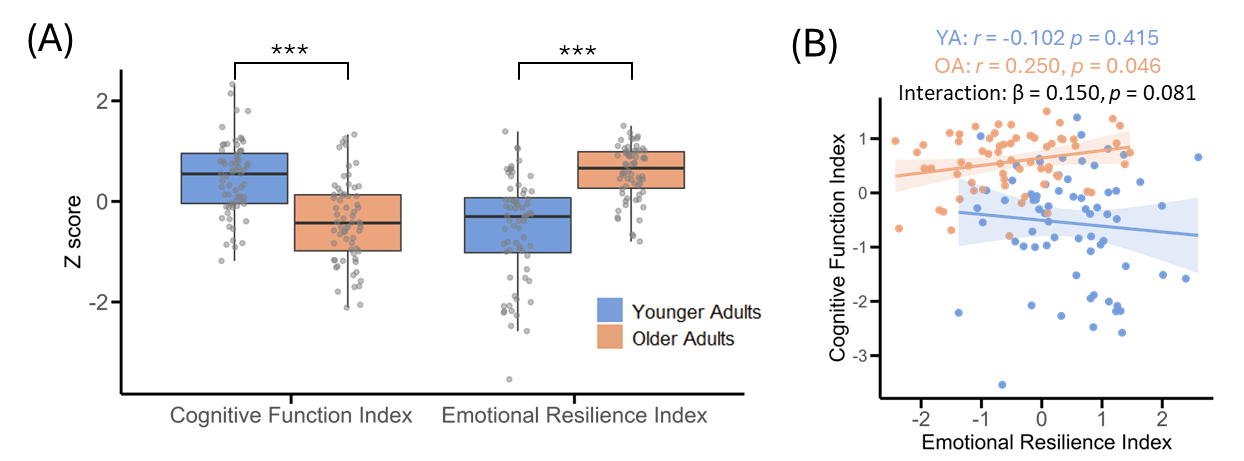

**Figure S3. Group differences and associations between emotional resilience index and cognitive function index.** (A) Group comparisons on emotional resilience index and cognitive function index between younger and older adults. The central black line represents the median, the box edges indicate the 25th and 75th percentiles (interquartile range), and the whiskers denote the data range (B) Scatter plots illustrating the relationship between emotional resilience index and cognitive function index for each age group. The solid line represents the fitted regression line, and the shaded area indicates the 95% confidence interval of the fit. YA= younger adults; OA=older adults. ^***^: *p* < 0.001.

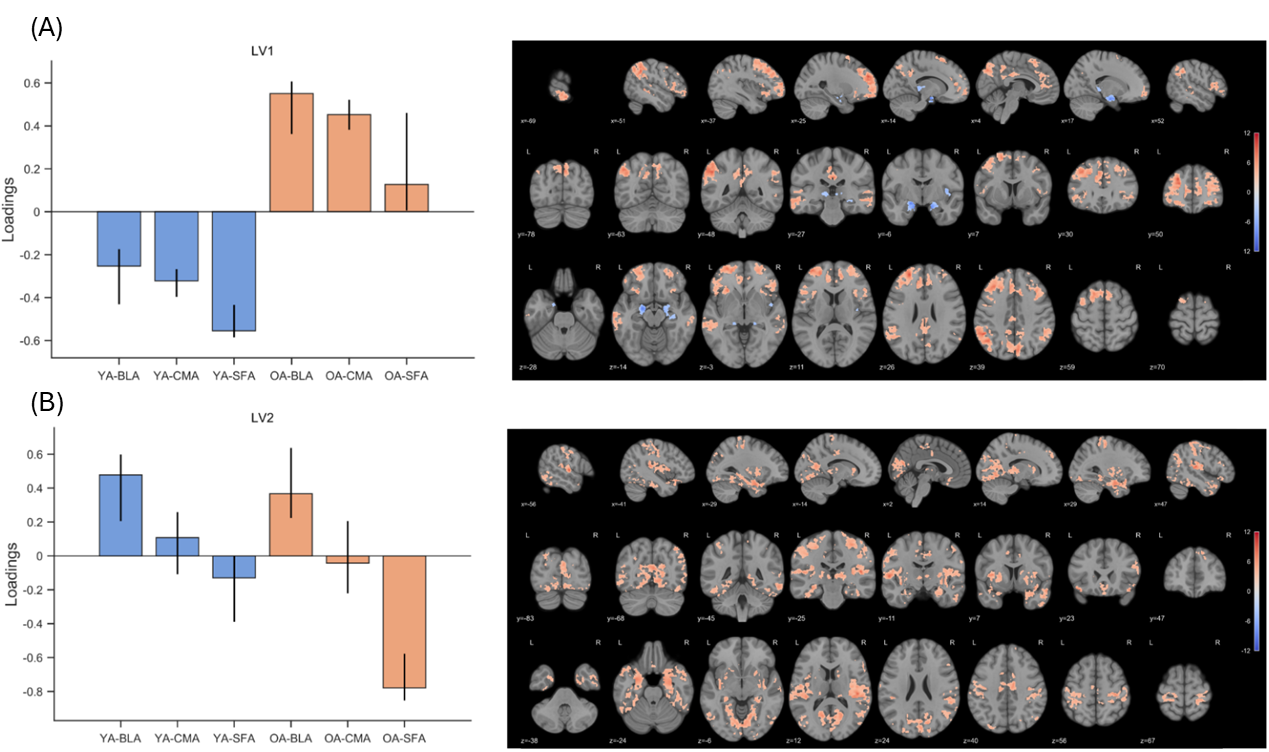

**Figure S4. Functional networks associated with amygdala subfields across age groups during the neutral movie. (A)** LV1 represents the overlap across all amygdala subfields engaged in older adults. The salience map shows areas engaged by older adults (yellow) and younger adults (blue), similarly across all subfields. **(B)** LV2 shows similar activation of the BLA in both younger and older adults. The brain salience map indicates regions that were positively connected with the BLA in both age groups. The map was thresholded at bootstrap ratios (BSR) ≥ |3.3| (approximately *p* < 0.001), with a minimum cluster size of 400 mm³ (≈205 voxels at 1.25-mm resolution).

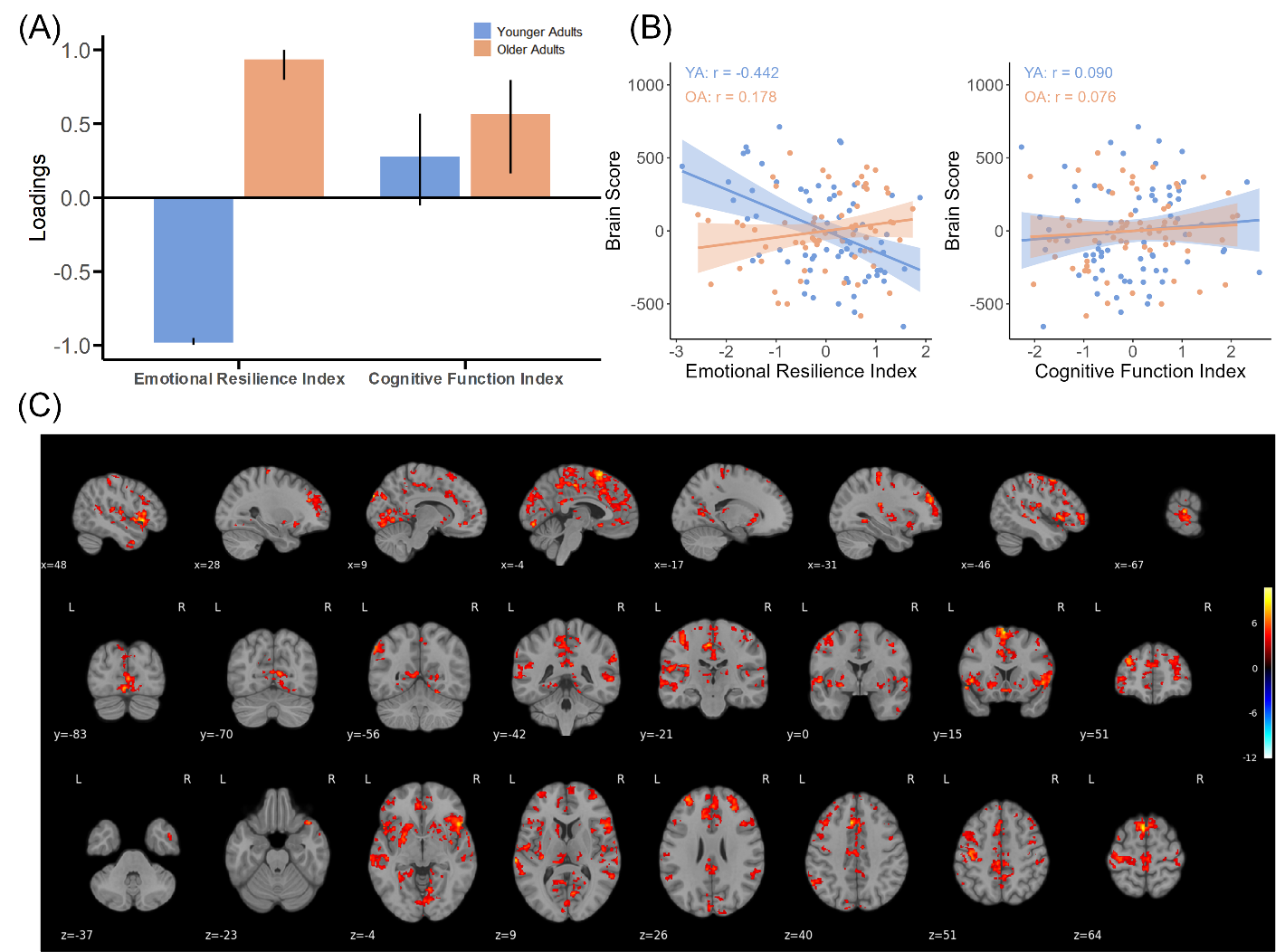

**Figure S5. PLS analysis of brain–behavior associations for LV1 of whole amygdala FC. (A)** The plot shows the behavioral loadings for each variable contributing to the significant latent variable (LV1). Error bars represent 95% confidence intervals derived from bootstrap resamples. Behavioral measures whose confidence intervals did not include zero were interpreted as reliably contributing to the identified brain–behavior pattern. **(B)** Scatterplots show the correlations between brain scores and the emotional resilience index and cognitive function index in two age groups. **(C)** Salience map showing the BLA-related functional network associated with behavioral measures. The map was thresholded at bootstrap ratios (BSR) ≥ |3.3| (approximately p < 0.001) with a minimum cluster size of 400 mm³ (≈205 voxels at 1.25-mm resolution).

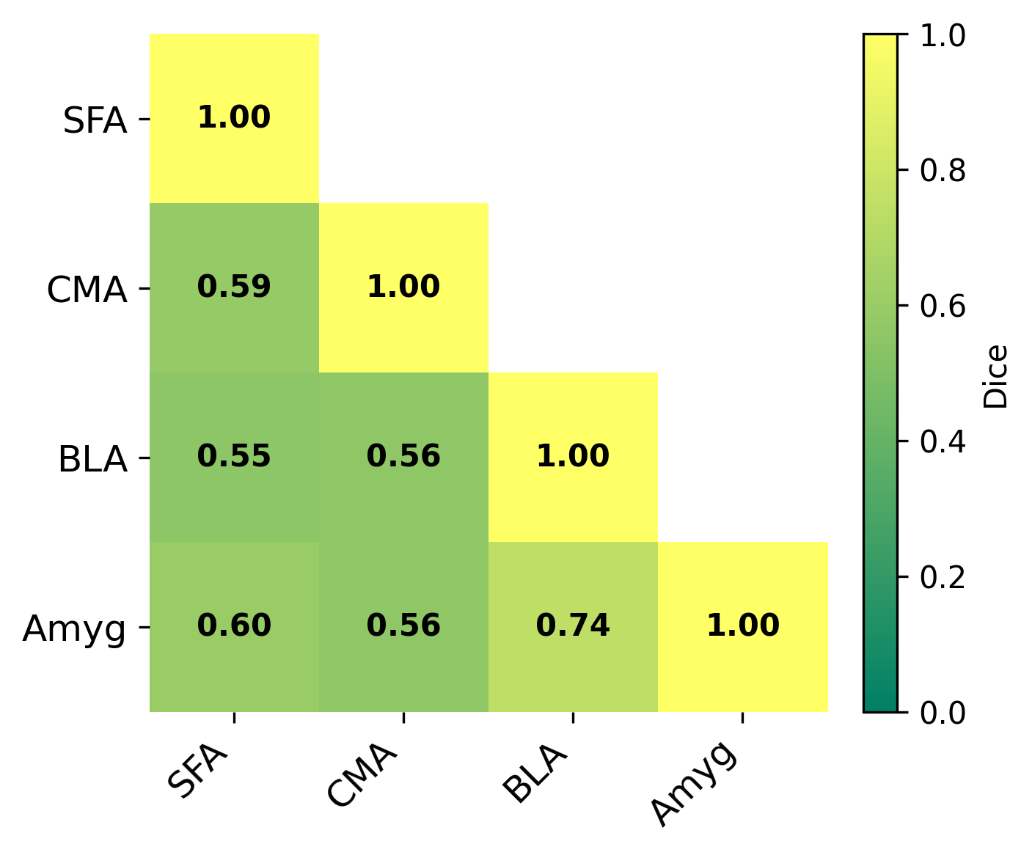

**Figure S6. Dice similarity coefficients between the salience maps of the amygdala and its subregions.** A higher Dice similarity coefficient indicates greater spatial correspondence between two functional patterns. BLA=basolateral amygdala; CMA=centromedial amygdala; SFA=superficial amygdala; Amyg=whole amygdala.
